# Stimulus-specific processing of the plasma membrane receptor-like kinase FERONIA in *Arabidopsis thaliana*

**DOI:** 10.1101/2021.10.08.463713

**Authors:** Cassidy S. Cornblatt, Han-Wei Shih, Gabriele B. Monshausen

## Abstract

FERONIA (FER), a receptor-like kinase involved in plant immunity, cell expansion, and mechanical signal transduction, is known to be endocytosed and degraded in response to treatment with its peptide ligand RAPID ALKALINIZATION FACTOR 1 (RALF1). Using confocal fluorescence microscopy and biochemical assays, we have found that full length FER-eGFP abundance at the plasma membrane is also regulated by mechanical stimulation, but through a separate, cysteine protease-dependent pathway. Like RALF1 treatment, both mechanical bending and mechanical wounding trigger a reduction in plasma membrane-localized, native promoter-driven FER-eGFP in Arabidopsis roots, hypocotyls, and cotyledons. However, pharmacological inhibition of protein trafficking and degradation suggests that while RALF1 induces clathrin-mediated endocytosis and subsequent degradation of FER-eGFP, mechanical stimulation triggers cleavage and/or degradation of FER-eGFP in a cysteine protease-dependent, clathrin-independent manner. Despite the stimulus-dependent differences in these two pathways, we found that both require early FER signaling components, including Ca^2+^ signaling, FER kinase activity, and the presence of LLG1, a FER-interacting protein with an essential role in FER-dependent signal transduction.

## INTRODUCTION

Receptor-like kinases (RLKs) play a central role in enabling plants to sense and respond to their biotic and abiotic environment. Multiple members of this superfamily bind to extracellular ligands, whereupon they initiate signal transduction cascades culminating in transcriptional and post-translational responses (Yin et al., 2019). In order to appropriately tune the duration and intensity of such responses, control mechanisms need to be in place to modulate RLK-mediated signaling. In animal systems, termination or reversible off-switching of receptor signaling can occur via receptor (de)phosphorylation, decreasing the availability of or affinity for the receptor ligand, proteolytic cleavage, and ubiquitination and endocytosis of receptors for recycling or eventual degradation (Ledda and Paratcha, 2007; Lemmon et al., 2016; Kliewer et al., 2017; Fernandes et al., 2019).

In Arabidopsis, plasma membrane levels of RLKs involved in defense signaling, including FLAGELLIN-SENSITIVE 2 (FLS2), PEP RECEPTOR 1/2 (PEPR1/2), EF-TU RECEPTOR (EFR), and LYSIN MOTIF-CONTAINING RECEPTOR-LIKE KINASE 5 (LYK5), have been shown to be controlled via ligand-induced endocytosis (Robatzek et al., 2006; Zipfel et al., 2006; Krol et al., 2010; Mbengue et al., 2016; Erwig et al., 2017). While many details about the mechanisms underlying activation and subsequent internalization of specific pattern recognition RLKs remain to be characterized, some common themes are emerging. The receptors heteromerize with co-receptors such as BRI1-ASSOCIATED RECEPTOR KINASE 1 (BAK1)/SOMATIC EMBRYOGENESIS RECEPTOR KINASE (SERK) family members or CHITIN ELICITOR RECEPTOR KINASE 1 (CERK1) upon binding their respective ligands (Wan et al., 2019), activation involves a phosphorylation step (Salamon and Robatzek, 2006; Macho et al., 2014; Ben Khaled, 2016; Erwig et al., 2017), and receptors appear to undergo ubiquitination before being endocytosed (Lu et al., 2011; Li et al., 2014a; Liao et al., 2017). Whether BAK1 and CERK1 co-receptors are internalized and degraded along with the associated pattern recognition RLKs is currently not known.

There is, however, intriguing evidence that BAK1 and CERK1 can be proteolytically processed directly at the plasma membrane (Petutschnig et al., 2014; Zhou et al., 2019). The proteolytic cleavage appears to be related to the RLKs’ role in cell death control, which is distinct from their role in immune responses (Schwessinger et al., 2011; Petutschnig et al., 2014; Zhou et al., 2019). This suggests that RLKs may be differentially processed for degradation in different signaling contexts. In animals, the receptor tyrosine kinase MET is a prototype for such diverging receptor processing pathways. Upon binding its ligand, the hepatocyte growth factor HGF, MET induces cell survival responses and undergoes endocytosis and lysosomal degradation. In the absence of its ligand, MET can be subjected to presenilin-regulated intramembrane proteolysis for removal under steady-state conditions, cleavage by caspases to enhance apoptotic signaling, or inactivation via calpain-mediated cleavage during necrosis (Fernandes et al., 2019).

In plants, FERONIA (FER), a member of the malectin-like-domain CrRLK1L subfamily, is involved in a wide range of growth, developmental, and stress response processes including cell expansion (Duan et al., 2010; Shih et al., 2014; Bastien et al., 2016), pollen reception (Huck et al., 2003; Rotman et al., 2003; Escobar-Restrepo et al., 2007), hormone responses (Deslauriers and Larsen, 2010; Yu et al., 2012; Höfte, 2015; Chen et al., 2016; Guo et al., 2018; Dong et al., 2019), defense signaling (Guo et al., 2018), salt stress responses (Yu and Assmann, 2018; Zhao et al., 2018), cell wall integrity maintenance (Feng et al., 2018; Dünser et al., 2019; Herger et al., 2019), and mechanical signaling (Shih et al., 2014). FERONIA has been proposed to serve as a scaffold to assemble immune receptors (Stegmann et al., 2017), but it is also known to function as a receptor for secreted cysteine-rich signaling peptides of the RAPID ALKALINIZATION FACTOR (RALF) family (Haruta et al., 2014; Zhao et al., 2018; Xiao et al., 2019; Gjetting et al., 2020; Yu et al., 2020). Binding of the RALF23 peptide appears to promote the heteromerization of FER with LORELEI-LIKE GLYCOSYLPHOSPHATIDYLINOSITOL-ANCHORED 1 and 2 (LLG1/2) *in vitro* (Xiao et al., 2019), and *in vivo*, LLG1 and LORELEI proteins are required to mediate FER-regulated signaling processes (Li et al., 2015; Galindo-Trigo et al., 2020).

Given its key role in integrating myriad physiological processes, the abundance of FER at the plasma membrane is likely to be tightly controlled. FERONIA was recently shown to be internalized in response to treatment with its RALF ligand (Zhao et al., 2018; Yu et al., 2020) and to also undergo endocytosis under steady state conditions (Yu et al., 2020). Here we present evidence that FER is processed via distinct degradation pathways in different signaling contexts. We confirm that RALF1 triggers clathrin-mediated endocytosis and degradation of FER along the well-established endosomal trafficking pathway. In contrast, mechanical stimulation by bending or wounding induces FER degradation via a cysteine protease-dependent process, which does not appear to involve FER internalization. While these divergent, stimulus-specific cellular fates of FER may reflect different functions of the receptor, both require operational FER signal transduction components.

## RESULTS

### Fungal RALF and mechanical stimulation elicit reduction of plasma membrane-localized FER-eGFP fluorescence similar to RALF1 treatment

Given the recent evidence that exogenous RAFL1, RALFL22, and RALF23 induce FER internalization (Zhao et al., 2018; Yu et al., 2020), we first wanted to confirm that FER plasma membrane (PM) abundance is also modulated by binding its RALF1 ligand under our experimental conditions. To this end, we used confocal microscopy to visualize PM FER-eGFP fluorescence in roots, hypocotyls, and cotyledons of 4- to 5-d-old transgenic *fer-4* mutants transformed with *FERpro:FER-eGFP*, a construct that fully complements *fer* null mutants (Shih et al., 2014). Fluorescence was quantified in ImageJ by outlining the PMs of three representative cells in each image and averaging their fluorescence intensities (Supplemental Figure S1). Since PM fluorescence intensity was variable in *FERpro:FER-eGFP/fer-4* seedlings, we calculated ratios (fluorescence_after treatment_/fluorescence_before treatment_) to normalize values. We found that RALF1 triggered a significant reduction in PM FER-eGFP fluorescence within 1 h of treatment in root and cotyledon epidermal cells (Figure 1). No equivalent loss of fluorescence was observed in control experiments, showing that reduction in fluorescence intensity was not due to photobleaching or mixing of growth medium (Figure 1, A, C-F). Together, these results confirm that RALF1 triggers FER internalization.

**Figure 1.**
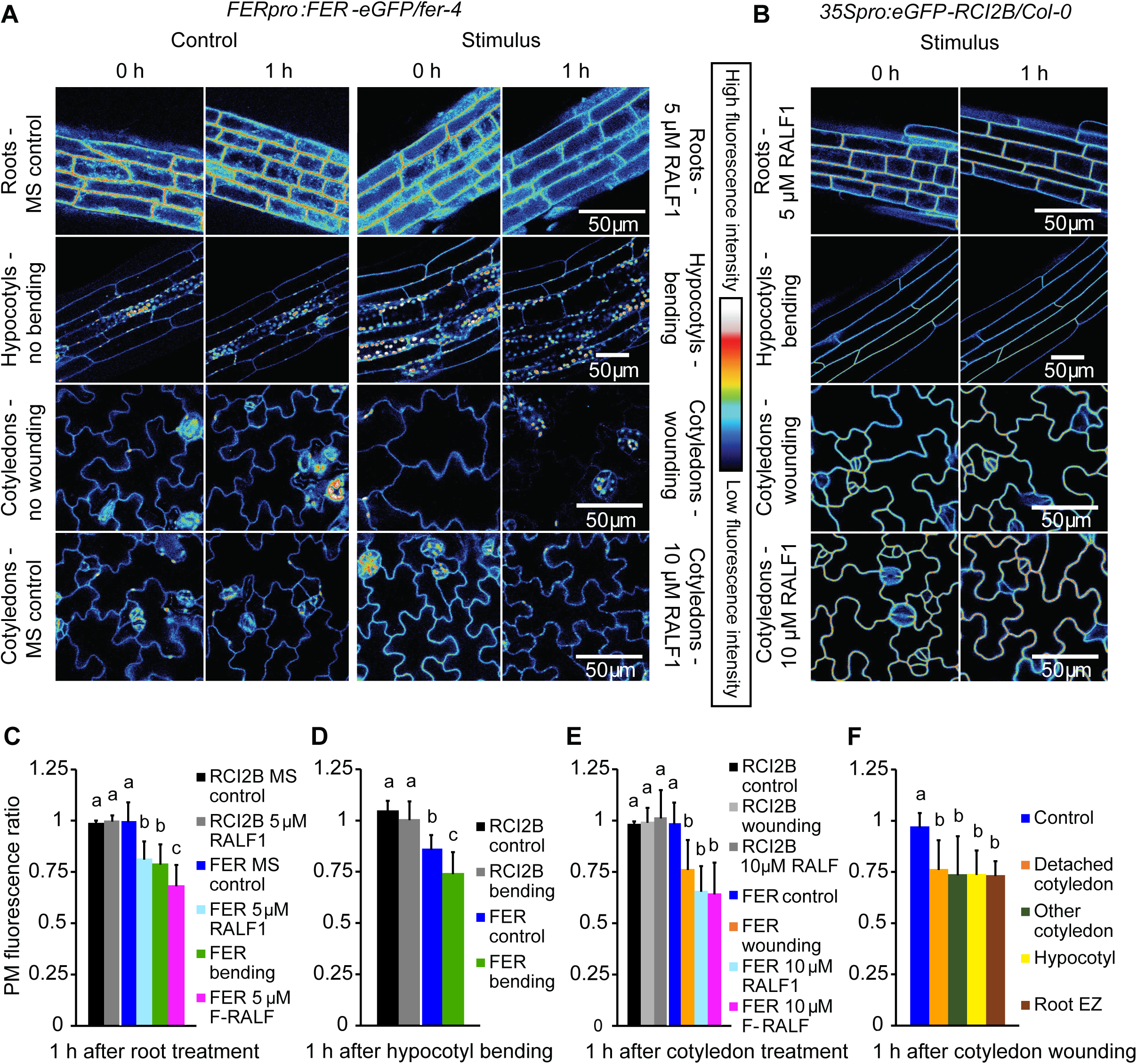
RALF1 and mechanical stimulation trigger a decrease in Arabidopsis FER-eGFP PM abundance. A, Fluorescence intensity of native promoter-driven FER-eGFP in epidermal PM decreases following RALF1 treatment of roots and cotyledons, bending of hypocotyls, or wounding by detaching cotyledons. Representative images of treated seedlings are shown alongside the respective controls. B, 35S-driven PM protein eGFP-RCI2B shows no decrease in fluorescence following stimulation. Representative images are shown. C-E, Quantification of PM fluorescence changes; ratios were calculated as PM fluorescence intensity 1 h after stimulation divided by fluorescence intensity before stimulation; values <1 thus indicate a decrease in fluorescence after treatment. RCI2B, *35Spro:eGFP-RCI2B/Col-0*; FER, *FERpro:FER-eGFP/fer-4*; F-RALF, *Fusarium*-derived RALF. C, Means + SD for n = 5 (RCI2B, MS control), 6 (RCI2B RALF1), 9 (FER MS control), 23 (FER RALF), 11 (FER bending), 8 (FER F-RALF) roots. D, Means + SD for n = 6 (RCI2B control), 6 (RCI2B bending), 7 (FER control), 7 (FER bending) hypocotyls. E, Means + SD for n = 4 (RCI2B control), 6 (RCI2B wounding), 6 (RCI2B RALF), 12 (FER control), 12 (FER wounding), 9 (FER RALF1), 8 (FER F-RALF) cotyledons. F, FER-eGFP PM fluorescence decreases throughout seedling in response to wounding by detaching a cotyledon. Means + SD of n = 12 (control), 12 (detached cotyledon), 6 (attached cotyledon), 4 (hypocotyl), 3 (root elongation zone) measurements. Letters indicate statistically significant differences (p ≤ 0.05; two-tailed Welch’s t-test).

What leads to increased release of endogenous RALF peptides to decrease FER PM abundance in plants is largely unknown; however, plants may be exposed to exogenous RALF when they encounter plant parasites or pathogens, some of which have been shown to secrete RALF peptide homologues during plant infection (Masachis et al., 2016; Thynne et al., 2017; Zhang et al., 2020). Indeed, FER-dependent RALF signaling appears to have been hijacked by some strains of the fungus *Fusarium oxysporum*, which may release F(*usarium*)-RALF to increase their virulence (Masachis et al., 2016; Thynne et al., 2017). To test whether F-RALF also triggers a decrease in FER-eGFP PM levels, we treated Arabidopsis roots and cotyledons with a synthetic RALF corresponding to the F-RALF produced by *F. oxysporum* f. sp. *lycopersici* 4287 (Thynne et al., 2017). We first confirmed that Arabidopsis roots produced a strong Ca^2+^ response to F-RALF similar to that observed after RALF1 treatment but with a more transient peak and a slight shoulder (Supplemental Figure S2). This difference in calcium signature is unsurprising since the kinetics of the Ca^2+^ responses to different Arabidopsis RALFLs are quite variable (Gjetting et al., 2020). We then examined the effect of F-RALF on FER-eGFP localization and found that F-RALF triggered a decrease in root and cotyledon epidermal cell PM FER-eGFP levels similar to RALF1 within 1 h of treatment (Figure 1, C and E). Fungal RALF thus appears to coopt not only FER-dependent ion signaling pathways but can also modulate the abundance of FER at the PM.

In addition to operating as a RALF peptide receptor, FER also serves a key role in cell wall integrity maintenance (Duan et al., 2010; Feng et al., 2018), although it is not yet clear if and how these functions are linked. As we have previously shown that FER is essential for normal mechanical signal transduction (Shih et al., 2014), we decided to analyze the effect of mechanical stimulation on FER-eGFP PM levels. Two types of mechanical stimulation were performed – (i) organ bending of roots and hypocotyls and (ii) mechanical wounding by detaching a cotyledon – and both types were found to elicit a decrease in PM FER-eGFP fluorescence (Figure 1, A, C-F; Supplemental Figure S3). Interestingly, the response to cotyledon wounding occurred throughout the seedling, including in both the detached and the still attached cotyledon as well as in the hypocotyl and the root elongation zone (Figure 1F).

Importantly, loss of FER-eGFP fluorescence in response to RALF1 treatment and mechanical stimulation does not appear to be part of an unspecific alteration of all PM protein levels as neither the plasma membrane marker eGFP-RCI2B/LTI6B (Cutler et al., 2000) nor FLS2-eGFP exhibited a reduction in fluorescence intensity under these conditions (Figure 1,B, C-E, and 2). Conversely, when we treated plants expressing FLS2-eGFP or FER-eGFP with the FLS2 ligand flg22, FLS2-eGFP PM fluorescence decreased as expected (Robatzek et al., 2006) while FER-eGFP fluorescence remained stable (Figure 2; see also Zhao et al., 2018). These results demonstrate the specificity of the RALF1- and mechanically-induced PM FER-eGFP reduction.

**Figure 2.**
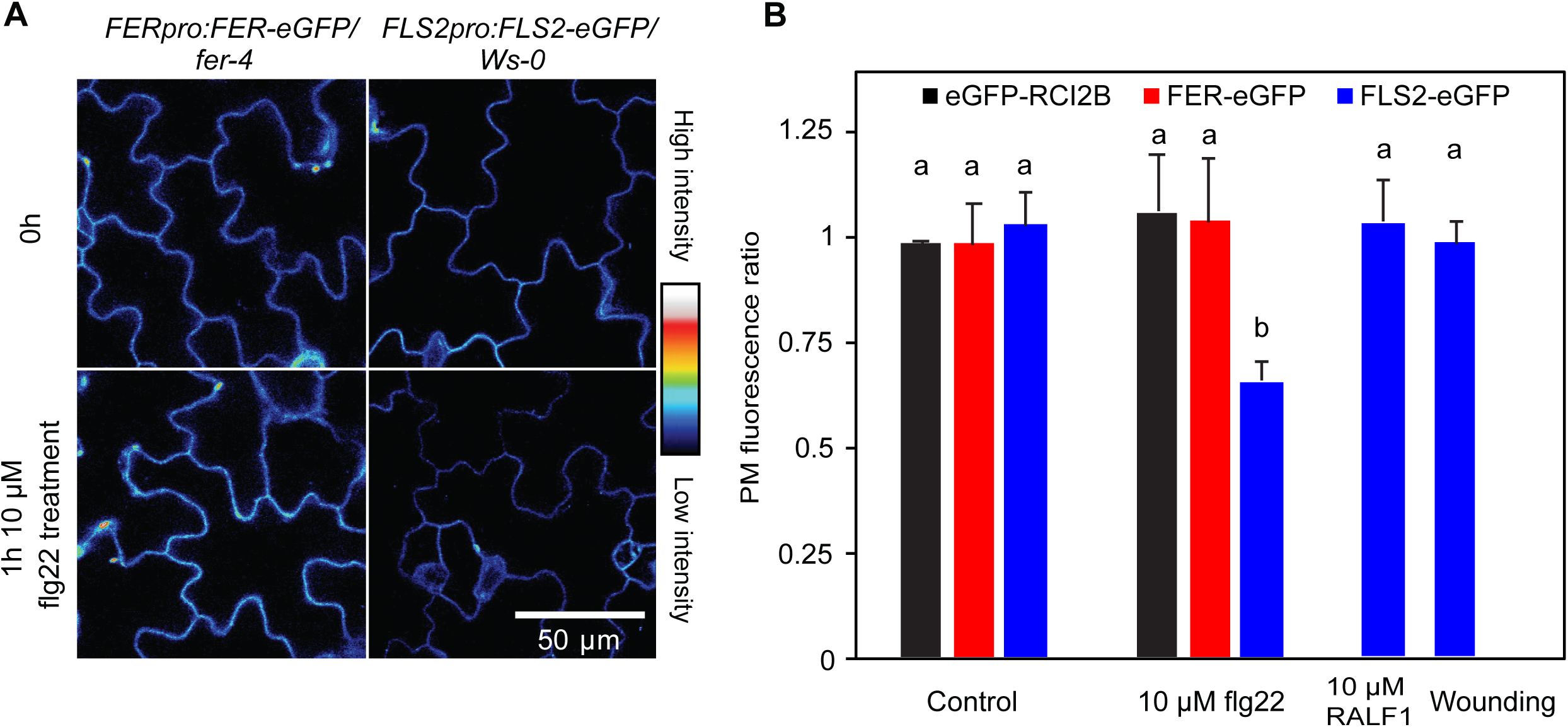
Loss of FER-eGFP and FLS2-eGFP PM fluorescence is stimulus-specific. A, Cotyledon fluorescence of native promoter-driven FER-eGFP and FLS2-eGFP before and 1 h after exposure to 10 µM flg22. B, Quantification of plasma membrane protein-eGFP fluorescence changes in response to different stimuli; ratios were calculated as PM fluorescence intensity 1 h after stimulation divided by fluorescence intensity before stimulation. Means + SD of n = 6 (FLS2 control), 4 (RCI2B flg22), 4 (FER flg22), 6 (FLS2 flg22), 4 (FLS2 RALF1), 4 (FLS2 wounding), seedlings; data for RCI2B and FER controls are taken from Figure 1. Letters indicate statistically significant differences (p ≤ 0.05; two-tailed Welch’s t-test).

Plasma membrane levels of FER-eGFP have previously been shown to be reduced by severe salt stress (Zhao et al., 2018). While we were able to confirm these results, we also observed a pronounced loss of eGFP-RCI2B fluorescence in seedlings treated with 150 mM NaCl (Supplemental Figure S4), indicating that the salt response is not specific to FER.

### Regulation of FER-eGFP PM abundance is dependent on Ca^2+^ signaling and requires an active FER kinase domain

Though it has previously been shown that FER undergoes internalization in response to RALF exposure, little is known about how the process is regulated. Since both RALF1 and mechanical perturbation trigger rapid, FER-dependent increases in cytosolic Ca^2+^ concentration ([Ca^2+^]_cyt_) in both Arabidopsis seedling roots and shoots (Haruta et al., 2008; Shih et al., 2014), and mechanical wounding is known to elicit a propagating wave of [Ca^2+^]_cyt_ throughout the plant (Toyota et al., 2018), we hypothesized that Ca^2+^ signaling functions upstream of stimulus-induced FER-eGFP removal from the PM. Previous aequorin-based measurements of RALF1-treated seedlings have not yet resolved which organs or tissues contribute to the Ca^2+^ response in shoots (Haruta et al., 2008), so we first monitored Arabidopsis expressing the genetically encoded Ca^2+^ sensor Yellow Cameleon 3.6 (Nagai et al., 2004) and confirmed that cotyledons indeed exhibit [Ca^2+^]_cyt_ elevation upon RALF1 exposure (Supplemental Figure S2B). Pretreatment with 300 µM of the Ca^2+^ channel blocker La^3+^ (Monshausen et al., 2009) abolished the RALF1- and mechanically-induced reduction of PM FER-eGFP fluorescence (Figure 3), indicating that this loss of PM FER-eGFP is dependent on Ca^2+^ signaling.

**Figure 3.**
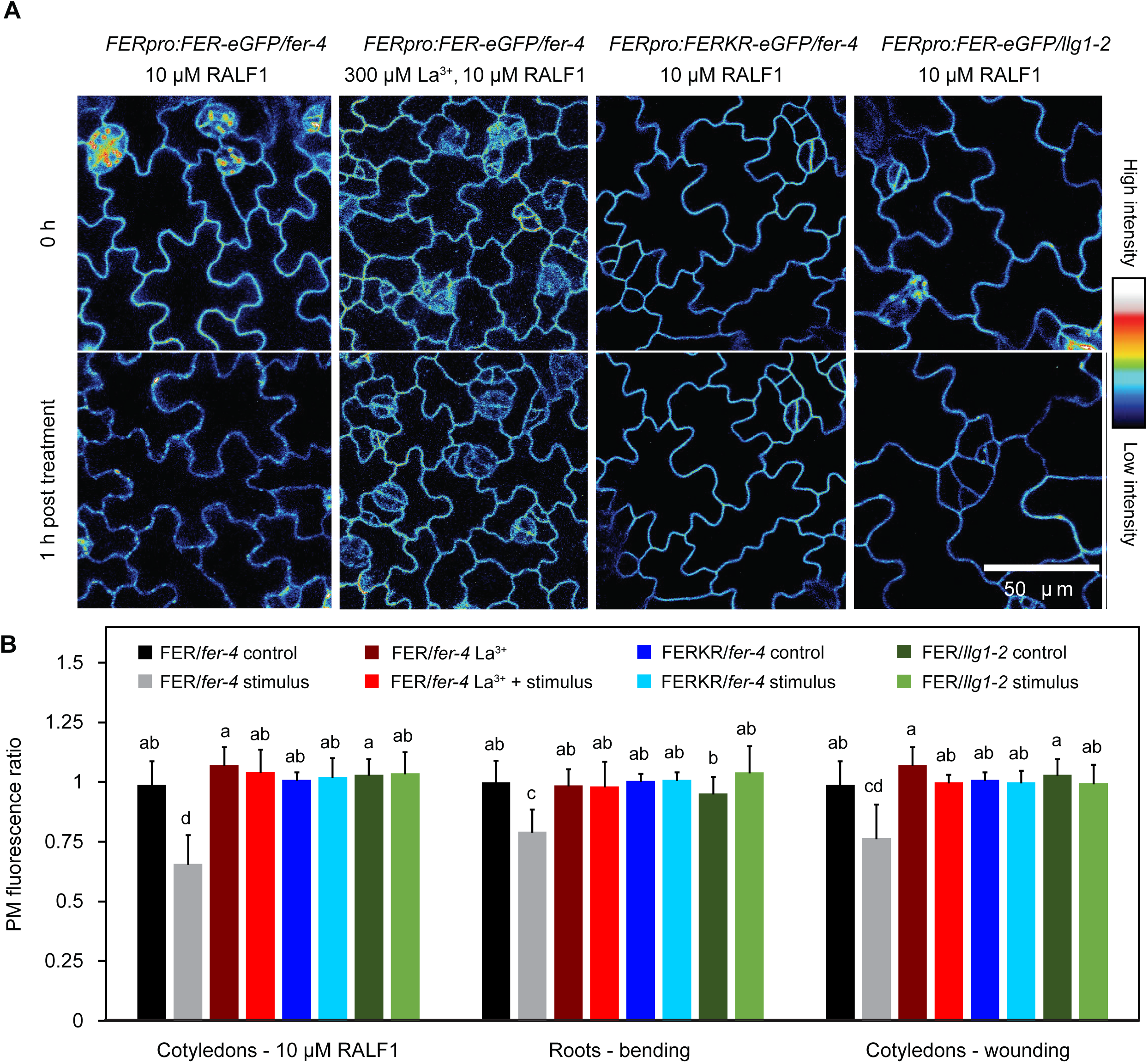
Stimulus-induced loss of FER-eGFP from the PM is dependent on Ca^2+^ signaling, FER kinase activity, and the presence of LLG1. A, Cotyledon fluorescence of Arabidopsis mutants expressing FER-GFP or kinase-inactive FERKR-GFP before and 1 h after treatment with 10 µM RALF1. To inhibit Ca^2+^ signaling, some seedlings were pretreated for 15 min with 300 µM La^3+^. B, Quantification of PM fluorescence changes in stimulated and corresponding control seedlings; ratios were calculated as PM fluorescence intensity 1 h after stimulation (or control) divided by fluorescence intensity before stimulation. Seedlings were stimulated either by treating with 10 µM RALF1, mechanical bending of roots, or wounding by detaching cotyledons. Means + SD for cotyledon RALF1 treatment: n = 5 (FER La^3+^), 6 (FER La^3+^ + RALF1), 7 (FERK control), 6 (FERK RALF1), 8 (FER/*llg1* control), 7 (FER/*llg1* RALF1) seedlings; for root bending assay: n = 12 (FER La^3+^), 12 (FER La^3+^ + bending), 7 (FERK control), 7 (FERK bending), 7 (FER/*llg1* control), 8 (FER/*llg1* bending); for cotyledon wounding assay: n = 5 (FER La^3+^), 5 (FER La^3+^ + wounding), 9 (FERK control), 8 (FERK wounding), 8 (FER/*llg1* control), 8 (FER/*llg1* wounding); data for FER-eGFP are taken from Figure 1. Letters indicate statistically significant differences (p ≤ 0.05; two-tailed Welch’s t-test).

FERONIA kinase activity is known to be involved in FER-dependent RALF1 and mechanical signaling but is not absolutely required for generation of Ca^2+^ and pH responses (Shih et al., 2014; Haruta et al., 2018) or Arabidopsis shoot development and stomatal movements (Chakravorty et al., 2018). We analyzed a *fer* null mutant expressing a kinase inactive version of FER (FERKR (K565>R) (Escobar-Restrepo et al., 2007); *FERpro:FERKR-eGFP/fer-4* line #7) and observed no decrease in FER-eGFP PM fluorescence following RALF1 treatment or mechanical stimulation (Figure 3), suggesting that the mechanism(s) controlling stimulus-induced changes in PM FER-eGFP abundance involve trans-and/or autophosphorylation by FER.

### LLG1 is required for normal FER-dependent responses and FER-eGFP endocytosis and degradation

FERONIA and RALF peptides have been shown to physically interact in a complex with the members of the GPI-anchored LRE/LLG protein family, which function as FER co-receptors (Li et al., 2015; Xiao et al., 2019). To determine whether LLG1 modulates FER-dependent ion signaling, we analyzed [Ca^2+^]_cyt_ and pH responses of *llg1* loss-of-function mutants in response to RALF1 and mechanical stimulation. In WT roots, RALF1 elicited a rapid increase in [Ca^2+^]_cyt_ and root surface pH as well as intracellular acidification, all of which were absent in *llg1* mutants (Figure 4) as well as *fer* mutants (Supplemental Figure S5; see also Haruta et al., 2008, 2014). Bending triggered a biphasic [Ca^2+^]_cyt_ increase in Arabidopsis wild-type (WT) roots, whereas two independent insertional mutants of LLG1, *llg1-1* and *llg1-2*, exhibited only the initial, FER-independent [Ca^2+^]_cyt_ elevation similar to *fer* mutants (Figure 4A; see also Shih et al., 2014). Furthermore, bending-induced root surface and cytosolic pH changes were only monophasic in *llg1* mutants (Figure 4, B and C), again phenocopying the *fer* mutant’s signaling defects (Supplemental Figure S5; see also Shih et al., 2014). The ion signaling response to RALF1 treatment and root bending was restored in *llg1* mutants complemented with *LLG1* driven by the *LLG1* native promoter (Supplemental Figure S5). These results confirm the essential role of LLG1 in FER-dependent ion signal transduction in roots.

**Figure 4.**
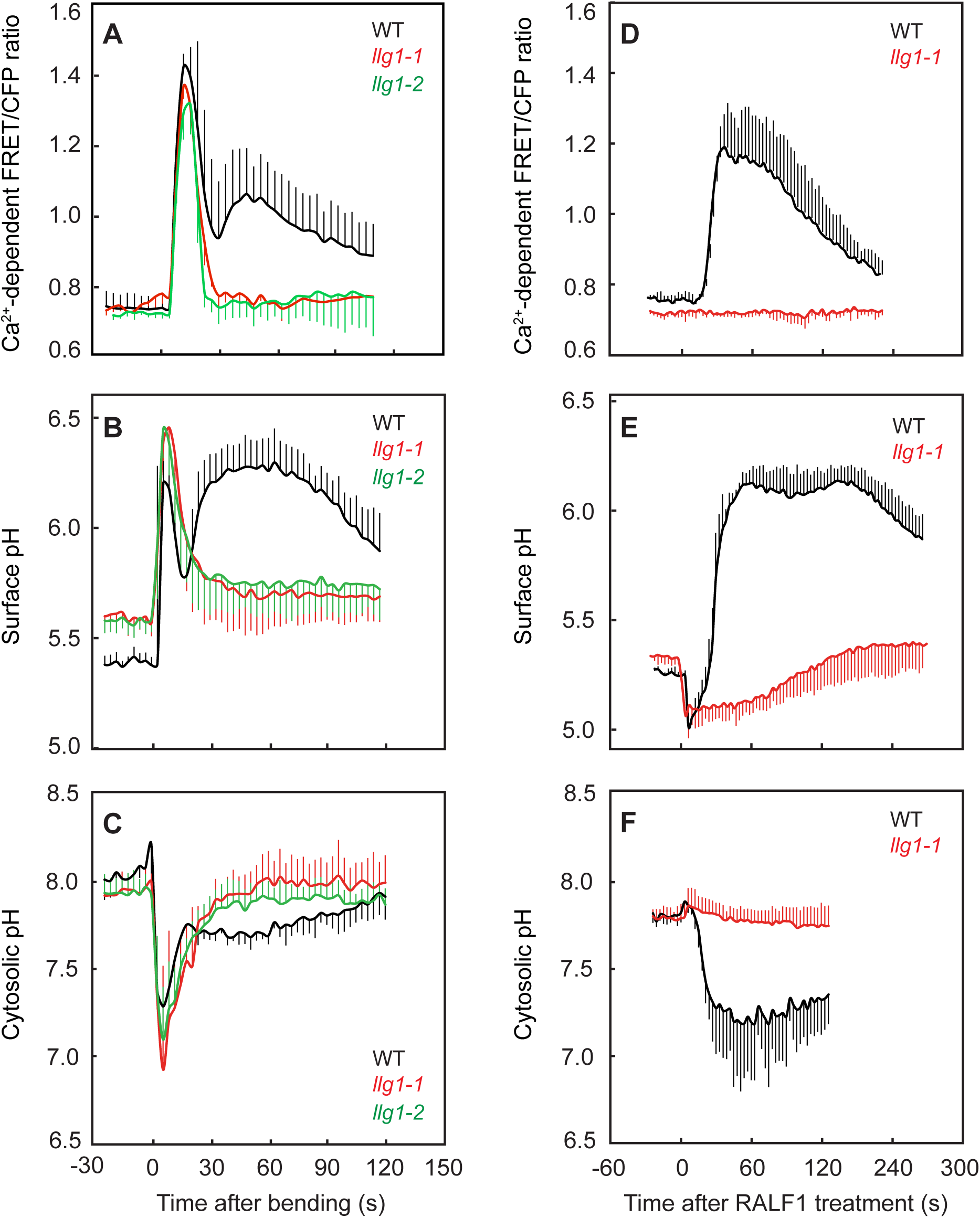
Arabidopsis *llg1* mutants exhibit *fer-*like ion signaling responses to root bending and RALF1. A-C, Effect of 1 µM RALF1 on (A) cytosolic Ca^2+^, (B) surface pH, and (C) cytosolic pH in epidermal cells of WT and *llg1* root elongation zone. Means + or – SD of n = (A) 4 (WT) and 5 (*llg1-1*); (B) 6 (WT), 7 (*llg1-1*); (C) 4 (WT), 5 (*llg1-1*) roots. D-F, Effect of root bending on (D) cytosolic Ca^2+^, (E) surface pH, and (F) cytosolic pH in epidermal cells on the convex (stretched) side of WT and *llg1* root elongation zone. Means + or – SD of n = (D) 3 (WT), 4 (*llg1-1*) 5 (*llg1-2*); (E) 6 (WT, *llg1-1 and llg1-2*); (F) 4 (WT, *llg1-1, llg1-2*) roots.

Consistent with an integral role in FER-dependent signaling pathways, *llg1* null mutants exhibited defects in cell expansion similar to *fer* mutants, including severe root hair defects, impaired control of diffuse cell expansion, and misshapen and bulging hypocotyl epidermal cells (Supplemental Figures S6 and S7). *Llg1* mutants also phenocopied *fer* mutants with regard to growth responses to external mechanical challenges, showing both pronounced right-handed skewing and abnormal barrier tracking responses (Supplemental Figure S8). Complementation of *llg1* restored normal mechanical development.

Given that *llg1* and *fer* mutants have virtually identical root signaling and developmental defects, we investigated whether mis-expression or mis-localization of FER in the *llg1* mutant background could account for *fer*-related phenotypes as previously proposed (Li et al., 2015; Shen et al., 2017). We introduced *FERpro:FER-eGFP* into the *llg1-2* mutant background and found that transgenic *llg1* seedlings showed the same FER-eGFP expression patterns and subcellular localization as *FERpro:FER-eGFP/fer-4* lines. In both *llg1* and *fer-4* backgrounds, FER-eGFP was clearly localized to the PM as well as intracellular punctae in epidermal cells of the root and cotyledon (Figure 3, Supplemental Figure S9). This suggests that loss of LLG1 abolishes FER-dependent mechanical signaling by affecting FER activity rather than FER trafficking to the PM.

LLG1 is also required for stimulus-dependent control of FER PM abundance. *Llg1-*2 mutants expressing *FERpro:FER-eGFP* showed no decrease in PM FER-eGFP fluorescence in epidermal cells of roots and cotyledons following either 1 h of RALF1 treatment or mechanical stimulation (Figure 3). Such an essential role for LLG1 in stimulus-induced modulation of FER-eGFP PM levels is consistent with its requirement for FER-dependent signaling and its physical interaction with FER in a heteromeric receptor complex (Xiao et al., 2019).

### Mechanically induced loss of PM FER-eGFP is insensitive to inhibitors of the endocytic/endosomal degradation pathway

Ligand-induced degradation of PM receptors is typically mediated by endocytosis and subsequent endosomal trafficking of the receptors to the vacuole. As we had not been able to detect an increase in FER-eGFP-labeled vesicles or endosomes after stimulation, we used a pharmacological approach to interrupt this pathway prior to cargo delivery to the vacuole. We first treated seedlings with 2 μM concanamycin A (ConcA), an inhibitor of the V-type H^+^-ATPase, to inhibit late endosome formation and thus degradation of internalized FER proteins (Huss et al., 2002; Ben Khaled et al., 2015). As anticipated, pre-incubation with ConcA triggered a more than sevenfold increase in the number of fluorescently labeled endosomes in RALF1-treated cotyledons (Figure 5). Conversely, no increase in labeled endosomes was detected in FERKR-eGFP-expressing seedlings pretreated with ConcA (Figure 5B), consistent with the lack of FERKR-eGFP internalization described above.

**Figure 5.**
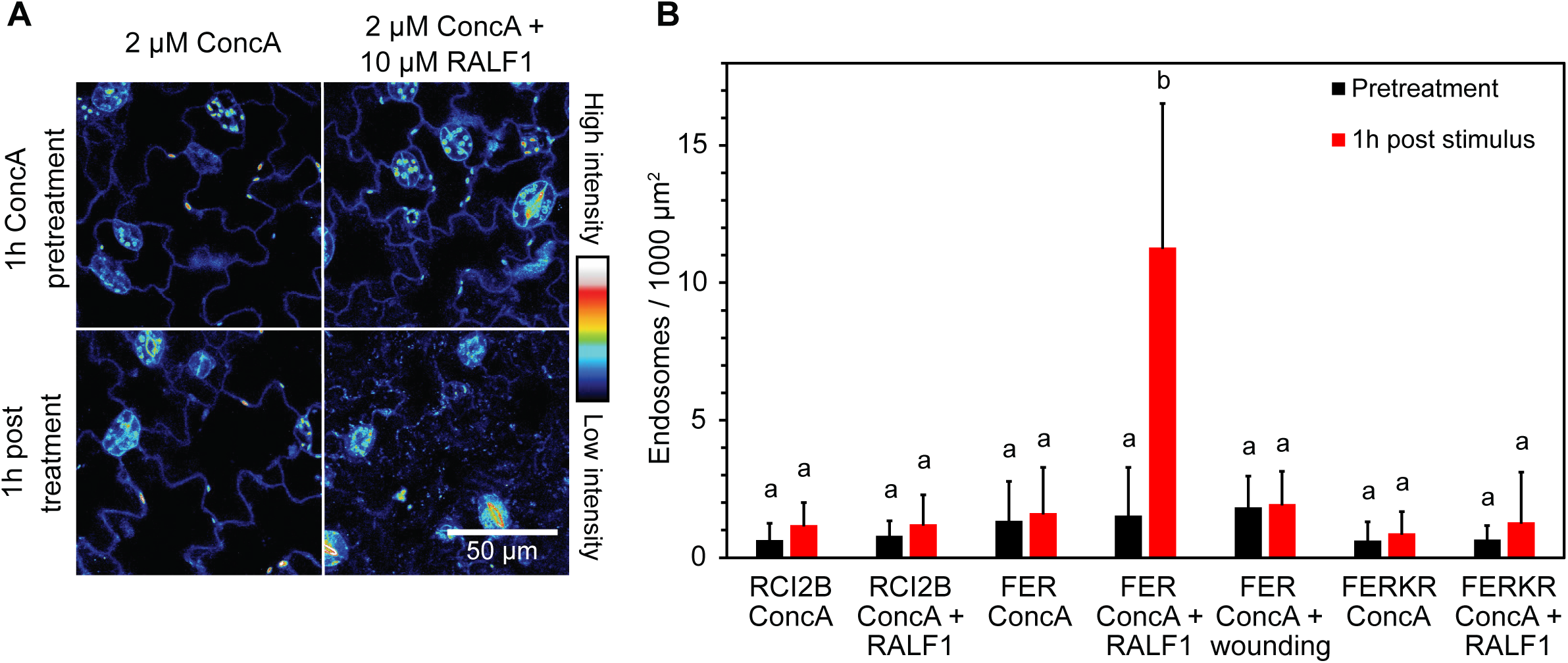
RALF1-induced loss of PM FER-eGFP is accompanied by appearance of FER-eGFP labeled endosomes in the presence of concanamycin A. A, Effect of ConcA on subcellular localization of FER-eGFP in pavement cells of Arabidopsis *FERpro:FER-eGFP/fer-4* cotyledons. Seedlings were first pretreated with 2 µM ConcA for 1 h and then incubated for 1 h with either 2 µM ConcA (control; left panels) or 2 µM ConcA + 10 µM RALF1 (right panels). Note the appearance of fluorescent punctae after RALF1 treatment, reflecting FER-eGFP-labeled endosomes. Representative fluorescence images are shown. B, Endosomal targeting is specific to RALF1-treatment and kinase-active FER-eGFP. Note that neither eGFP-RCI2B nor FERKR-eGFP appeared in endosomes after treatment with 2 µM ConcA + 10 µM RALF1. Mechanical wounding of cotyledons also triggered no detectable internalization of FER-eGFP. Endosomal density was quantified by segmentation analysis. Means + SD of n = 5 (RCI2B ConcA), 5 (RCI2B ConcA + RALF1), 8 (FER ConcA), 8 (FER ConcA + RALF1), 6 (FER ConcA + wounding), 5 (FERKR ConcA) and 5 (FERKR ConcA + RALF1) seedlings. Black bars, number of endosomes after 1h ConcA pretreatment; red bars, number of endosomes after 1h treatment (only ConcA or ConcA + stimulus). Letters indicate statistically significant differences between each pretreatment and corresponding post treatment (p ≤ 0.05; two-tailed Welch’s t-test).

Surprisingly, the number of fluorescently labeled intracellular punctae also did not increase in ConcA-pretreated cotyledons after wounding, suggesting that the decrease in FER-eGFP PM fluorescence observed in response to mechanical perturbation does not involve the endosomal degradation pathway (Figure 5B).

To further verify that RALF1 and wounding trigger FER degradation via different pathways, we incubated seedlings with 30 µM wortmannin (Wm), which is known to inhibit clathrin-mediated endocytosis (Emans et al., 2002; Takagi and Uemura, 2018). As expected, preincubation with Wm for 30 min blocked the disappearance of PM-localized FER-eGFP upon RALF1 treatment (Figure 6). Wortmannin-pretreated seedlings still exhibited a normal RALF1-induced [Ca^2+^]_cyt_ increase, indicating that Wm did not generally reduce sensitivity to RALF1 (Supplemental Figure S2C). In contrast, Wm pretreatment did not impair the wounding-induced loss of PM FER-eGFP fluorescence (Figure 6, A and C).

**Figure 6.**
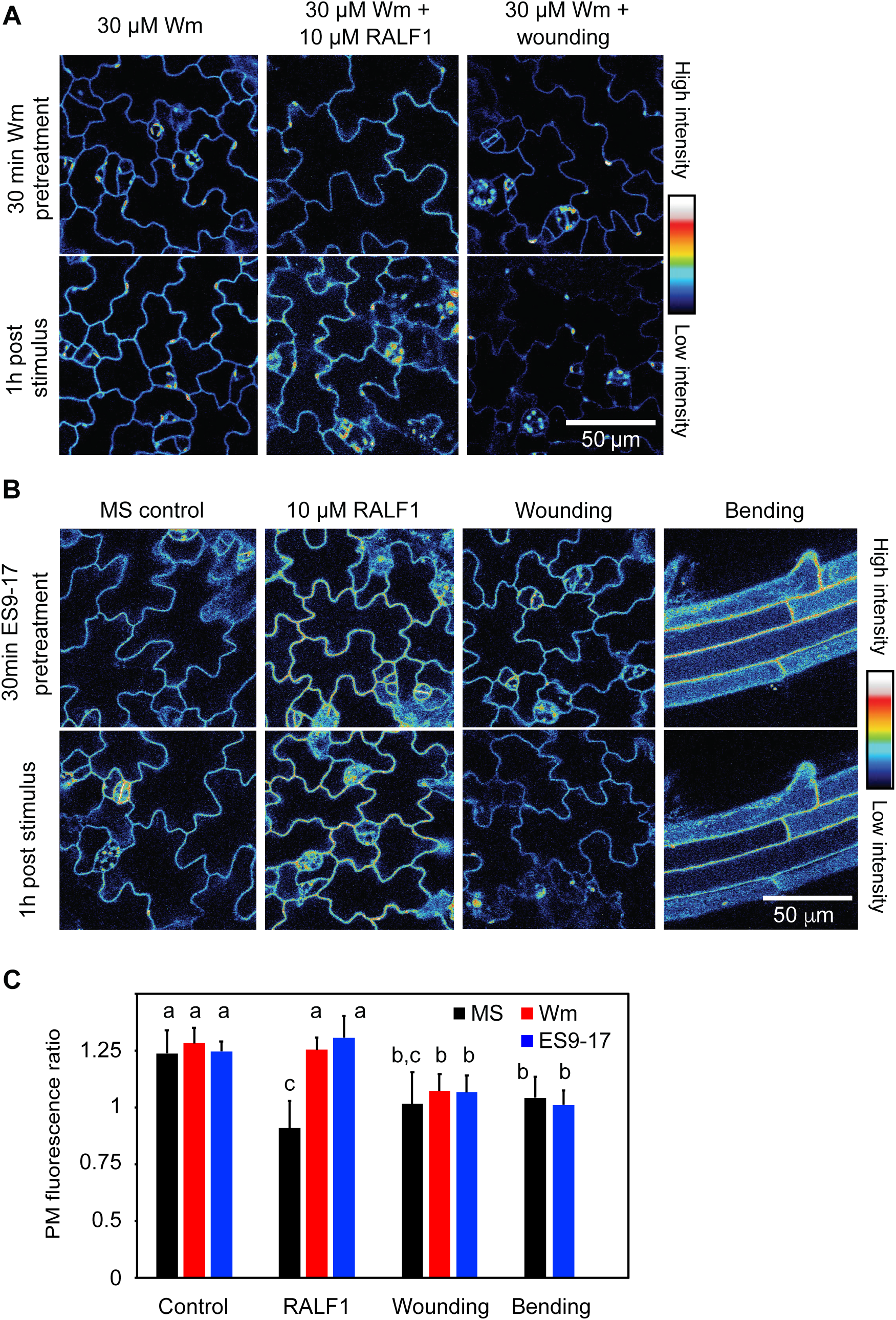
Differential effects of wortmannin and Endosisin 9-17 on RALF1- and mechanically-induced loss of FER-eGFP PM fluorescence. A, Effect of Wm on FER-eGFP PM abundance in pavement cells of Arabidopsis pFER::FER-eGFP/fer-4 cotyledons. Seedlings were pretreated with 30 μM Wm for 30 min and then incubated for 1 h with either 30 μM Wm (control; left panels) or 30 μM Wm + 10 µM RALF1 (middle panels), or subjected to wounding in the presence of 30 μM Wm (right panels). Representative fluorescence images are shown. B, Effect of ES9-17 on FER-eGFP PM abundance in epidermal cells of Arabidopsis FERpro:FER-eGFP/fer-4 seedlings. Seedlings were pretreated with 30 μM ES9-17 for 30 min and then incubated for 1 h with either 30 μM ES9-17 (control) or 30 μM ES9-17 + 10 µM RALF1, or subjected to mechanical stimulation by cotyledon wounding or root bending in the presence of 30 μM ES9-17. Representative fluorescence images are shown. C, Quantification of FER-eGFP PM fluorescence changes; ratios were calculated as PM fluorescence intensity 1 h after stimulation divided by fluorescence intensity before stimulation. Means + SD of n = 4 (Wm), 8 (Wm + RALF1), 8 (Wm + wounding), 5 (ES9-17), 6 (ES9-17 + RALF1), 4 (ES9-17 + wounding), 8 (ES9-17 + root bending) seedlings; data for FER-eGFP MS are taken from Fig. 1. Letters indicate statistically significant differences (p ≤ 0.05; two-tailed Welch’s t-test).

These results were further corroborated when we blocked clathrin-mediated endocytosis using the recently developed Endosidin 9-17 (ES9-17), a specific inhibitor of clathrin heavy chain function (Dejonghe et al., 2019). Pretreatment with 30 µM ES9-17 for 30 min abolished the RALF1-induced reduction in FER-eGFP PM fluorescence, while ES9-17 had no effect on either the wounding- or bending-induced reduction in FER-eGFP PM fluorescence (Figure 6, B and C).

Taken together, these results provide evidence that the decrease in PM FER-eGFP fluorescence in response to mechanical stimulation occurs via a pathway that is distinct from the previously established clathrin-mediated, RALF1-induced FER endocytosis pathway (Yu et al., 2020).

### Mechanical stimulation triggers cysteine protease-dependent degradation of FER-eGFP

To determine whether FER-eGFP is targeted for degradation upon mechanical stimulation, we treated seedlings with MG132, an inhibitor commonly used to block proteasomal degradation by the 26S proteasome (Lee and Goldberg, 1998). Preincubation with 50 µM MG132 for 30 min completely abolished the reduction in FER-eGFP PM fluorescence upon mechanical wounding and bending but did not affect the RALF1 response (Figure 7, Supplemental Figure S10). Surprisingly, lactacystin, a more specific inhibitor of the 26S proteasome (Fenteany et al., 1995), did not inhibit the wounding-induced reduction of the FER-eGFP PM fluorescence, suggesting that the effect of MG132 was independent of the 26S proteasome (Figure 7, A and B, Supplemental Figure S10).

**Figure 7.**
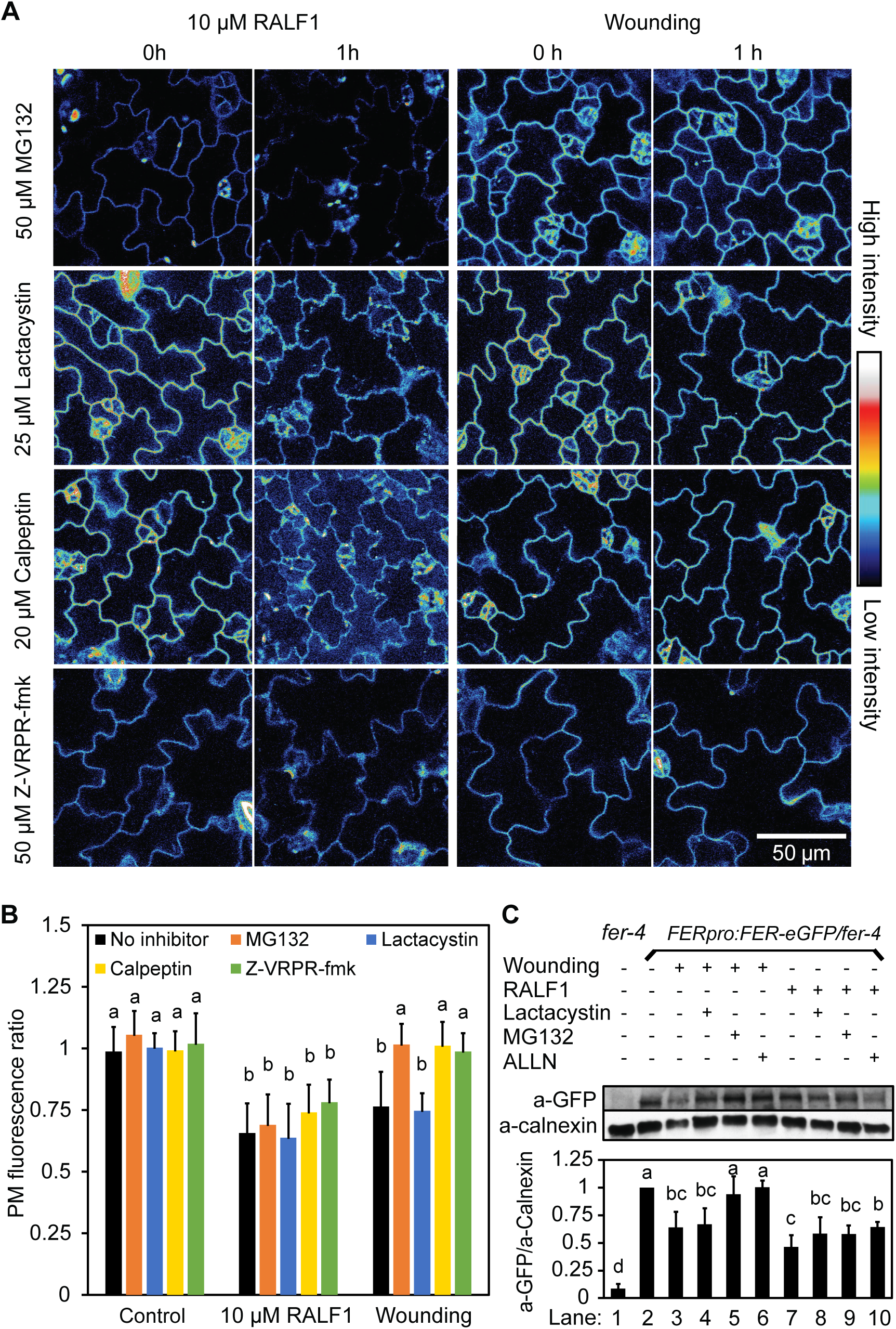
Differential effects of protease inhibitors on RALF1- and wounding-induced FER-eGFP degradation. A, Effect of protease inhibitors on FER-eGFP PM abundance in pavement cells of Arabidopsis *FERpro:FER-eGFP/fer-4* cotyledons. Seedlings were pretreated for 30 min with either the 26S-proteasome inhibitor lactacystin, the general cysteine protease inhibitor MG132, the calpain inhibitor calpeptin or the metacaspase inhibitor Z-VRPR-fmk and then either incubated for 1 h with protease inhibitor + 10 µM RALF1 or subjected to wounding in the presence of the protease inhibitor. Representative fluorescence images before and after stimulation are shown. B, Quantification of FER-eGFP PM fluorescence changes; ratios were calculated as PM fluorescence intensity 1 h after stimulation divided by fluorescence intensity before stimulation. Means + SD for control: n = 5 (MG132), 4 (lactacystin), 4 (calpeptin), 5 (Z-VRPR-fmk); for RALF1 treatment: n = 10 (MG132), 4 (lactacystin), 4 (calpeptin), 5 (Z-VRPR-fmk); for wounding assay: n = 10 (MG132), 7 (lactacystin), 4 (calpeptin), 5 (Z-VRPR-fmk) seedlings; data for FER-eGFP ‘no inhibitor’ are taken from Figure 1. C, Immunoblot analysis of total protein extracted from Arabidopsis *FERpro:FER-eGFP/fer-4* seedling shoots 3 h after treatment by RALF1 exposure or wounding. The immunoblots were probed with antibodies against GFP and calnexin as indicated, with the latter used as a loading control. Blot quantification was performed by calculating the ratios of anti-GFP band density values by corresponding density values of anti-calnexin bands (see also Supplemental Figure S12). Note that pretreatment with the cysteine protease inhibitor MG132 and the calpain inhibitor ALLN inhibited wounding-induced, but not RALF1-induced FER-eGFP degradation. The shown immunoblot is representative of three biological replicates. Means + SD of n = 4 blots except for ALNN treatment (n=3); band density ratios were normalized to the untreated control (lane 2). Letters indicate statistically significant differences (p ≤ 0.05; two-tailed Welch’s t-test).

Unlike lactacystin, MG132 is known to also inhibit the activity of several cysteine proteases, including calpains (Tsubuki et al., 1996). We therefore decided to further investigate the involvement of these cysteine proteases in mechanically-induced loss of PM FER-eGFP fluorescence. Intriguingly, two different calpain inhibitors, calpeptin and ALLN (Ono et al., 2016), and two different metacaspase inhibitors, MALT1 Inhibitor 1 (MI-2) and Z-VRPR-fmk (Hander et al., 2019), completely abolished the decrease in PM FER-eGFP abundance specifically after mechanical stimulation (Figure 7, A and B, Supplemental Figures S10 and S11), suggesting that the loss of PM fluorescence may be the result of FER-eGFP degradation or cleavage by cysteine proteases in the calpain and/or metacaspase families.

To verify that loss of PM fluorescence was accompanied by a reduction in FER-eGFP protein levels, we immunoblotted protein extracted from seedlings stimulated by RALF1/wounding treatment in the absence or presence of protease inhibitors (Figure 7C, Supplemental Figure S12). While we did not detect any fragment that would correspond to a product of C-terminal FER-eGFP cleavage, the Western blots did show a clear reduction in full-length FER-eGFP levels upon wounding and RALF1 exposure. In agreement with our imaging data, the wounding-induced FER-eGFP decrease was inhibited by MG132 and ALLN but not the 26S proteasome inhibitor lactacystin, while the RALF1-induced decrease was not sensitive to any of these drugs, indicating that FER undergoes differential, stimulus-dependent processing.

## DISCUSSION

In this paper, we have shown that PM localization of the RLK FER is dynamically regulated not only by RALF1, but also by other stimuli known to activate FER-dependent signaling cascades. Exposure to Arabidopsis RALF1, a fungal (*Fusarium oxysporum*)-RALF, mechanical bending of roots and hypocotyls, and mechanical wounding of cotyledons led to a 20-40% reduction in PM FER-eGFP fluorescence within 1 h of the start of each treatment (Figure 1). These treatments did not affect the abundance of the PM marker RCI2B/LTI6B or the flg22 receptor FLS2 (Figures 1 and 2), indicating that the responses are at least partially specific to FER. Given that inhibition of Ca^2+^ signaling, disruption of FER kinase activity, and lack of the FER co-receptor LLG1 all abolished the stimulus-induced loss of PM FER-eGFP, the response further seems to be conditioned on an operational FER signal transduction machinery (Figure 3).

FERONIA PM abundance appears to be post-translationally controlled via distinct mechanisms. Some environmental conditions, such as salt stress, trigger the internalization of assorted PM proteins, including FER, PIN1 and PIN2, BRI1, PIP2;1, and RCI2B/LTI6B (Li et al., 2011; Zwiewka et al., 2015; Zhao et al., 2018) (Supplemental Figure S4). This is likely a general cellular mechanism to quickly adjust cell surface area to a shrinking volume and is not dependent on activation of FER signaling pathways (Supplemental Figure S4), consistent with a role for FER in the recovery phase to salt stress rather than early signaling events (Feng et al., 2018). In contrast, FER-eGFP was at least somewhat selectively removed from the PM upon RALF1-treatment and mechanical stress (Figures 1 and 2), i.e., in response to activation of signal transduction pathways where FER plays an essential role in the earliest phases of signaling (Haruta et al., 2014; Shih et al., 2014). Moreover, FER-eGFP removal was insensitive to treatment with the FLS2 ligand flg22 (see also Zhao et al., 2018), even though FER is known to interact with FLS2 and has been shown to modulate FLS2-BAK1 receptor complex formation (Stegmann et al., 2017).

Surprisingly, given the shared requirement for early signaling components such as the presence of the co-receptor LLG1, elevation of cytosolic Ca^2+^ concentration and auto/transphosphorylation, the specific FER degradation pathways activated by mechanical stimulation and RALF1 appear to diverge downstream of these signaling events. As recently shown (Yu et al., 2020) and confirmed in our pharmacological assays, RALF1 elicits FER degradation via the well-established clathrin-mediated endocytosis/endosomal pathway previously described for PAMP-activated RLKs (Paez Valencia et al., 2016). In seedlings pretreated with the V-type H^+^ ATPase inhibitor ConcA, RALF1-induced loss of FER-eGFP from the PM was associated with its appearance in intracellular punctae (Figure 5), and both Wm and ES9-17 blocked the PM signal reduction (Figure 6), while inhibitors of proteasomal degradation or cysteine proteases had no effect (Figure 7).

Conversely, none of the inhibitors of the endocytic/endosomal pathway interfered with loss of PM FER-eGFP fluorescence induced by mechanical stimulation (Figures 5 and 6). Given the insensitivity to ConcA, we conclude that mechanically-induced degradation does not proceed via the endosomal or selective autophagy pathways, as both pathways are known to be disrupted by the inhibition of the V-type H^+^ ATPase (Bassham et al., 2006; Dettmer et al., 2006; Yang et al., 2019). Instead, mechanically-induced FER-eGFP degradation was highly sensitive to cysteine protease inhibitors (Figure 7). The *Arabidopsis thaliana* genome encodes a large number of cysteine proteases. Among these, cytosolic type II metacaspases are attractive candidates as they have been shown to cleave tonoplast-localized PROPEP precursors in a Ca^2+^-dependent manner in response to wounding and flg22 treatment (Hander et al., 2019; Shen et al., 2019). Another attractive candidate is DEK1, the only plant calpain, which has been implicated in mechanical signal transduction. DEK1 consists of a cytosolic calpain domain linked to a large membrane spanning region (Lid et al., 2002). The transmembrane domain appears to function either as a mechanically activated Ca^2+^ channel or as a mechanosensitive regulator of a Ca^2+^ channel (Tran et al., 2017), whereas DEK1 calpain activity is enhanced by Ca^2+^ (Wang et al., 2003), consistent with our observation that mechanically-induced FER-eGFP degradation requires Ca^2+^ signaling (Figure 3). Whether a type II metacaspase and/or DEK1 are indeed involved in the degradation of FER-eGFP remains to be determined.

How precisely FER-eGFP is processed at the plasma membrane is currently not known. Unlike the recent report of cysteine-protease inhibitor-sensitive cleavage of BAK1 (Zhou et al., 2019), we did not detect a C-terminal fragment that would result from the hydrolysis of a single peptide bond (simple cleavage); FER-eGFP may thus have been completely degraded. Alternatively, if the unknown cysteine protease(s) only had access to the cytoplasmic domain of FER, it is conceivable that an N-terminal FER fragment remained associated with the plasma membrane or was released into the cell wall space and retained some function. Such proteolytic cleavage has been reported for multiple animal tyrosine receptor kinases (Huang, 2021), but has also been observed in plants for the chitin-binding LysM-RLKs CERK1 and LYK4 under control conditions (i.e., in the absence of any particular stimulus) (Petutschnig et al., 2014). In the latter examples, the ectodomains of CERK1 and LYK4 were shed, while the CERK1 kinase domain likely remained associated with the plasma membrane. Determining whether FER ectodomains also persist after wounding, either as soluble fragments or bound to the membrane, will be key to understanding whether different cellular fates of FER have functional significance for the diverse signaling pathways FER is involved in.

One of the most interesting questions emerging from our investigation is how different stimuli can activate different degradation pathways for the same receptor. Differential phosphorylation of FER is an obvious candidate mechanism as FER kinase activity is required for degradation in response to both mechanical and RALF1 stimuli (Figure 3). Indeed, distinct FER phosphorylation patterns have already been detected in phosphoproteomic analyses under different experimental conditions. For example, FER phosphorylation has been detected at amino acids S^858^, S^871^, and S^874^ upon RALF1 treatment (Haruta et al., 2014), while S^695^, T^696^, and S^701^ were phosphorylated after exposure to flg22 (Nühse et al., 2004). Such treatment-dependent differential phosphorylation may mark FER for distinct cell fates. Some exciting recent work has revealed the importance of the phosphocode for receptor activity (Perraki et al., 2018; Latorraca et al., 2020), and at least for the well-studied receptor tyrosine kinase MET, it has been shown that the presence of receptor phosphorylation at a juxtamembrane tyrosin recruits a ubiquitin ligase, leading to internalization and lysosomal degradation, and impedes caspase-mediated proteolytic cleavage of MET (Fernandes et al., 2019). It remains to be determined if FER residues are phosphorylated in response to mechanical wounding and if the resulting phosphorylation pattern is specific to this signal transduction pathway.

Another important question relates to the functional significance of the divergent cellular fates of FER. Receptor signaling post internalization has been demonstrated in animals, where, for instance, MET was found to be capable of modulating the actin cytoskeleton through different pathways depending on its endosomal localization (Ménard et al., 2014). Post-internalization signaling has also been suggested to occur in response to RALF1 treatment, as RALF1-triggered root growth inhibition was found to be reduced in mutants defective in clathrin-mediated endocytosis (Yu et al., 2020).

RALF-regulated abundance of FER at the plasma membrane may also be an important element determining the virulence of fungal pathogens. Given that *Fusarium oxysporum* F-RALF was equally effective at triggering a decrease in PM FER-eGFP as endogenous RALF1 (Figure 2), we speculate that fungi are coopting RALF1-induced FER endocytosis and degradation in order to inhibit FER-supported defense signaling. Indeed, (Masachis et al., 2016) found that immune responses of tomato roots were enhanced when infected with a mutant *F. oxysporum* strain lacking the *F-RALF*-encoding gene, suggesting that F-RALF of WT *F. oxysporum* plays a role in suppressing host immunity. Interestingly, (Thynne et al., 2017) observed no significant difference in the virulence of *f-ralf* knockout *F. oxysporum* strains. Our data may explain these contradictory results. Unlike Masachis et al., Thynne et al. first wounded the plant roots prior to inoculating them with the mutant *F. oxysporum*, which may have triggered proteolytic processing of tomato FER well before exposure to the fungus, possibly preventing F-RALF from having an effect on virulence.

If N-terminal FER cleavage products remain associated with the plasma membrane, these could retain some non-kinase-dependent activity. FERONIA is proposed to function as a scaffold to facilitate interaction of immune receptors EFR and FLS2 with their BAK1 co-receptor (Stegmann et al., 2017), and it is possible that a truncated FER continues to regulate receptor-complex formation. Interestingly, it was recently demonstrated that FER kinase activity is not required for its role in immune signaling (Gronnier et al., 2020); whether the presence of the intracellular kinase domain is necessary remains to be seen.

In conclusion, we have shown that the abundance of full-length FER at the plasma membrane is regulated in a stimulus-dependent manner via distinct processes, that nevertheless all require FER signaling competency. Whether there are separate pools of FER targeted for differential processing by different signal transduction pathways, which post-translational modifications underlie the different cellular fates of FER, and what the functional significance is of these different fates are exciting questions for future research.

## METHODS

### Plant material and growth conditions

*Arabidopsis thaliana* seeds were surface sterilized with 50% bleach and plated on 1% agar containing ¼-strength Murashige and Skoog salts (MP Biomedicals) and 1% sucrose at pH 5.8. They were stratified for 2 d at 4ºC and grown for 4-5 d at 22ºC under continuous light. Seeds of the Arabidopsis T-DNA insertion mutants *fer-4* (GABI_106A06), and *llg1-1* (SAIL_47_G04), *llg1-2* (SALK_086036) were obtained from the Arabidopsis Biological Research Center (ABRC). *35Spro:eGFP-RCI2B* seeds (Cutler et al., 2000) were obtained from ABRC courtesy of Dr. Marisa Otegui (University of Wisconsin, Madison). Seeds of transgenic Arabidopsis expressing *FLS2pro:FLS2-3xMyc-eGFP* (Robatzek et al., 2006) were obtained from Dr. Antje Heese (University of Missouri) with kind permission of Dr. Silke Robatzek (Ludwig Maximilian University of Munich).

### Plasmid construction and plant transformation

*Llg1* loss-of-function mutants were complemented with *LLG1pro:LLG1-GFP* constructs. The promoter region was identified with the help of the Arabidopsis Gene Regulatory Information Server (AGRIS) resource. The constructs were cloned by first amplifying LLG1pro:LLG1 from genomic DNA using a forward primer annealing to the 5’-end of the predicted LLG1 promoter region and containing a SalI restriction site (5’-AGgtcgacGAAGTTTCGGTTTATATTTGATTATCTAA-3’) and a reverse primer directed to the 3’-end of the LLG1 coding sequence and containing a NotI restriction site (5’-TTgcggccgcGAGAACAACTTAACAAAAA-3’). The LLG1pro:LLG1 fragment was subcloned into the Gateway pENTR1a vector (Life Technologies) and recombined into the modified Gateway-compatible destination vector pEarleyGate302 (Earley et al., 2006). Constructs were introduced into *Agrobacterium tumefacien* strain GV3101 and transformed into Arabidopsis plants via floral infiltration. *FERpro:FER-eGFP* and *FERpro:FERKR-eGFP* Arabidopsis lines were created as described by Shih et al. (2014).

### FER-GFP fluorescence assays

#### Fluorescence microscopy

Fluorescence of GFP-tagged FER, RCI2B, and FLS2 was monitored using a Zeiss LSM 510 META confocal microscope and a 40x 1.2 NA C-Apochromat water immersion objection. eGFP was excited with the 488 nm line of an argon laser, and emission was collected with a 488 nm beamsplitter and a 505-550 nm band pass emission filter. Eight bit, 512 px × 512 px images were acquired with scan times of 1.97s and 2× line averaging.

Images were processed in ImageJ (http://rsbweb.nih.gov/ij/), and the data were analyzed with Microsoft Excel. Plasma membrane fluorescence was quantified by carefully outlining the plasma membrane as shown in Supplemental Figure 1 and measuring the average signal intensity. Three representative, in-focus cells in each image for each seedling were selected and averaged. Ratios of fluorescence intensities (post treatment/pretreatment) were used to normalize the data as different seedlings expressing the same construct exhibited quite variable baseline fluorescence intensities. Gray scale images of FER-eGFP fluorescence were transoformed into pseudo-color using the Royal LUT in ImageJ for better visualization of fluorescence intensities.

Endosomes were quantified as described by Leslie & Heese (2017). In brief, segmentation analysis was performed in ImageJ to select the endosomes in an automated fashion without selecting the plasma membrane or other fluorescent particles, and the number of endosomes was counted. Note that while we previously had to use a lower excitation laser intensity for the FERKR line to avoid signal saturation and could thus have missed a reduction in fluorescence intensity if the decrease was of the same absolute magnitude as observed for FER-eGFP, for endosomal analysis we imaged cotyledons using the same settings as for FER-eGFP (i.e., settings under which the PM signal was saturated).

#### RALF treatment

Seedlings were treated either with synthetic RALF1, corresponding to the mature Arabidopsis RALF1 (ATTKYISYQSLKRNSVPCSRRGASYYNCQNGAQANPYSRGCSKIARCRS; Biomatik) or synthetic fungal RALF, corresponding to the RALF produced by *Fusarium oxysporum* f. sp. *lycopersici* 4287 (SGEISYGALNRDHIPCSVKGASAANCRPGAEANPYNRGCNAIEKCRGGV; GenScript; Thynne et al., 2017).

For root imaging assays, 4- to 5-d-old Arabidopsis seedlings expressing GFP-tagged FER, RCI2B, and FLS2 were transferred to experimental chambers and embedded in 1% agar containing ¼-strength Murashige and Skoog (MS) salts and 1% sucrose, pH 5.8. Seedlings were kept in these chambers overnight in a vertical orientation to let them acclimate and to recover from mechanical perturbation during transfer (for a detailed description, see Bhat et al., 2021). Fluorescence in epidermal cells of the root elongation zone was imaged before and 1 h after agar was carefully cut away from the root tips and 5 μM RALF (dissolved in ¼ MS and 1% sucrose, pH 5.8) was added. Incubation with ¼ MS with 1% sucrose at pH 5.8 without RALF was used as a control.

For cotyledon imaging assays, 4- to 5-d-old Arabidopsis seedlings were selected based on detectable fluorescence in cotyledon pavement cells. Each selected seedling was fully immersed for 1 h in a 0.2 mL PCR tube containing either 10 μM RALF in ¼ MS and 1% sucrose, pH 5.8, or the control solution, ¼ MS and 1% sucrose, pH 5.8, without RALF. Cotyledons of each seedling were imaged before and 1 h after treatment.

#### Mechanical stimulation

For bending assays, 4-d-old Arabidopsis seedlings were transferred to experimental chambers and embedded in 2% agar containing ¼ MS and 1% sucrose, pH 5.8. Seedlings were kept in these chambers overnight in a vertical orientation. During the experiment, the roots or hypocotyls were gently bent approximately 90° with forceps for about 2 s and released. Because it was difficult to keep exactly the same cells in the field of view before and after bending, cells in the bent region were imaged immediately after and then again 1 h after bending. Unbent seedlings served as a control.

For wounding assays, 4- to 5-d-old Arabidopsis seedlings were selected based on detectable fluorescence in cotyledon pavement cells. One cotyledon from each selected seedling was detached just above the petiole using scissors and placed in ¼ MS and 1% sucrose, pH 5.8, alongside the wounded seedling. Fluorescence was imaged before and 1 h after wounding. All data is for the detached cotyledon unless otherwise noted.

#### flg22 treatment

Four-to five-day-old Arabidopsis seedlings with detectable fluorescence in cotyledon pavement cells were fully immersed for 1 h in Eppendorf tubes containing either 10 μM flg22 (Alpha Diagnostic International, Inc.) dissolved in ¼ MS and 1% sucrose, pH 5.8, or the control solution, ¼ MS and 1% sucrose, pH 5.8, without flg22. One cotyledon of each seedling was imaged before and after treatment.

#### Inhibitor experiments

The protocol followed was the same as described above for RALF1 and mechanical stimulation, but seedlings were pre-incubated with one of the following inhibitors for the indicated amount of time: 2 μM concanamycin A (1 h pre-incubation; BioViotica), 30 μM wortmannin (30 min pre-incubation; LC Laboratories), 30 μM endosidin 9-17 (30 min pre-incubation; Carbosynth Ltd.), 50 μM MG132 (30 min pre-incubation; Enzo), 25 μM lactacystin (30 min pre-incubation; AdipoGen), 20 μM calpeptin (30 min pre-incubation; Sigma-Aldrich), 20 μM ALLN (30 min pre-incubation; MilliporeSigma), 50 μM Z-VRPR-fmk (30 min pre-incubation; Enzo), 10 μM MALT1 Inhibitor I (30 min pre-incubation; EMD Millipore Corp), and 300 μM LaCl_3_ (in phosphate free ¼ MS and 1% sucrose; 15 min pre-incubation; Acros).

### Ion imaging

Monitoring of root surface and cytosolic pH changes in response to RALF1 or mechanical bending was performed essentially as previously described by (Monshausen et al., 2009, 2011). In brief, Arabidopsis seedlings were embedded in agar containing 10% Hoagland medium and 1% sucrose, ∼pH 5.1. After a recovery period of 4-6 h, the agar was cut away from the root tips and replaced with liquid medium. For cytosolic pH measurements, the medium consisted of 10% Hoagland medium and 1% sucrose, ∼pH 5.1; for root surface pH measurements the medium was additionally supplemented with 150 µg ml^-1^ of the pH-sensitive fluorophore fluorescein conjugated to 10 kDa dextran (Sigma-Aldrich). Seedlings were then either bent by 60-90° using a glass capillary or treated with an equal volume of medium containing 10 or 20 µM RALF peptides, for a final concentration of 5 or 10 µM. To measure cytosolic pH, the pH probe GFP H148D (35Spro:GFP H148D in pEarleyGate100 (Earley et al., 2006; Monshausen et al., 2007)) was stably introduced into Arabidopsis WT and mutants *fer-2, llg1-1* and *llg1-2*. Ratiometric imaging GFP H148D and fluorescein-dextran was performed using an inverted Zeiss LSM 510 META confocal microscope and a 10X 0.3NA Plan-Neofluar objective, with line-alternating excitation at 458 nm and 488 nm, a 488 nm dichroic mirror, and 505 nm long pass emission filter. Fluorescence ratios were calibrated using a series of buffers. For calibration of extracellular pH, buffers calibrated to pH 5.0 (100 mM malic acid), pH 5.5 (100 mM succinic acid), pH 6.0 (100 mM MES), pH 6.5 (100 mM PIPES), pH 7.0 (100 mM MOPS) and pH 7.5 (100 mM HEPES) were supplemented with fluorescein-dextran; for intracellular pH measurements, an endpoint calibration was performed using 100 mM NH_4_Cl, pH 9.3, and 100 mM KHCO_3_, pH 6.2 (see (Monshausen et al., 2007)).

To image [Ca^2+^]_cyt_, Arabidopsis WT, *llg1-1*, and *llg1-2* seedlings stably overexpressing Yellow Cameleon 3.6 (35Spro:YC3.6 in pEarleyGate100 (Monshausen et al., 2007)) were transferred to custom-made chambers and embedded in agar containing ¼ Murashige and Skoog medium and 1% sucrose, pH 5.7. The agar was then cut away from the region of interest and replaced with a solution containing ¼ Murashige and Skoog medium and 1% sucrose before treatment with RALF1, F-RALF1, or bending by 60-90° using a glass capillary. The sensor was excited at 458 nm, and Ca^2+^-dependent CFP and FRET emission were collected at 484-505 nm and 526-536 nm, respectively.

Ion signaling was monitored in/along epidermal cells of the root elongation/mature zone and in pavement cells in cotyledons. Image J was used to calculate the ratios of pH- and Ca^2+^-dependent fluorescence intensities in selected regions of interest.

### Western blotting

Approximately 20 5-d-old seedlings were transferred from agar plates to Eppendorf tubes containing ¼ MS, 1% sucrose, pH 5.8, and any indicated inhibitors. Following any necessary preincubation, seedlings were treated with 10 μM RALF1 or wounded and let sit for 3 h, after which roots were removed immediately prior to freezing seedling shoots with liquid nitrogen. The aerial tissue was ground up on ice using a plastic pestle, and protein was extracted as described by Li et al., (2014) except as noted using a buffer consisting of 10 mM Tris/Cl pH 7.5, 150 mM NaCl, 0.5 mM EDTA, 1% NP-40 (VWR Life Science), 1 mM PMSF (GoldBio), 1X Halt Protease Inhibitor Cocktail (Thermo Scientific), and 1 mM DTT (GoldBio). Protein concentration was measured using the Bradford method (Bradford, 1976), and samples were diluted to equivalent concentrations with extraction buffer. Samples of about 120-160 μg were loaded into wells of a 10% Mini-PROTEAN TGX Precast Protein Gel (Bio-Rad Laboratories) and run for 2.5 h at 90 V in buffer consisting of 25 mM Tris, 192 mM glycine, and 0.1% SDS, pH 8.3. The transfer was conducted on ice for 1 h at 100 V in buffer consisting of 25 mM Tris, 192 mM glycine, and 10% methanol. Antibody staining employed PBST buffer (from PBS 10X powder concentrate, Fisher BioReagents), and 10% milk was used for blocking. Gels were incubated with primary antibodies for 2 h and with horse-radish peroxidase (HRP)-conjugated secondary antibodies for 1 h. Antibody dilutions used were 1:500 for α-GFP (B-2, Santa Cruz Biotechnology), 1:2500 for α-calnexin (CNX1/2, Agrisera), 1:1000 for goat-α-mouse-HRP (AP124P, EMD Millipore), and 1:10000 for goat-α-rabbit-HRP (AS09 602, Agrisera). Both (FER)GFP and calnexin were visualized with the HRP substrate SuperSignal West Femto Maximum Sensitivity (Thermo Scientific) using a Konica SRX-101A.

Western blot bands were quantified as shown in Supplemental Figure 12c-d. Lanes were selected in ImageJ and analyzed using the Gel Analyzer function. Blot density was calculated, and the density for the α-GFP band was divided by the corresponding α-calnexin band density to calculate relative band density.

### Root skewing assay

Arabidopsis seeds were surface sterilized and sown on 1.5% agar containing ½ Murashige and Skoog salts and 1.5% sucrose, pH 5.7, and stratified at 4°C for 2 d. The plates were placed vertically for 3 d at 21°C under 16 h long-day photoperiod and tilted to 45° for another 3 d. Plates were scanned using the Epson Perfection V700 scanner, and root skewing angles were measured using Image J.

### Root barrier assay

Four-day-old Arabidopsis seedlings were transferred to experimental chambers and embedded in 0.7%, 2%, or 3% agar containing ¼ Murashinge and Skoog salts and 1% sucrose, pH 5.7. Twelve hours later, a coverglass sliver was inserted into the agar perpendicular to the root long axis close to the root tip (Shih et al., 2014) and the chamber was placed onto the stage of a Zeiss Axioplan microscope flipped on its back for vertical stage microscopy. Images of roots were acquired using a Zeiss 5X 0.25 NA Fluar objective and Stingray camera (Allied Vision Technologies), and root angles were measured in ImageJ.

### *Kinematic analysis of* Arabidopsis *root growth*

Four-day-old seedlings were transferred to experimental chambers and embedded in 1% agar containing ¼ Murashinge and Skoog salts and 1% sucrose, pH 5.7. After an overnight recovery period, the experimental chamber was mounted on a ZEISS Axioplan light microscope in vertical orientation. Images of growing roots were acquired every 30 s for 40 min using a Zeiss 10X 0.25NA Achroplan objective and Stingray F-504B monochrome camera (Allied Vision; Exton, PA, USA). Root relative elemental growth rate profiles and root elongation zone width were calculated as described previously by Shih et al., (2014).

Sequence data from this article can be found in the GenBank/EMBL data libraries under soybean gene accession numbers: FER (AT3G51550), LLG1 (AT5G56170), FLS2 (AT5G46330), RCI2B (AT3G05890).

## Supplemental Data

The following materials are available in the online version of this article.

Supplemental Figure S1. Ca^2+^ responses to Arabidopsis and *Fusarium* RALF peptides.

Supplemental Figure S2. Quantification of FER-eGFP PM fluorescence.

Supplemental Figure S3. Root bending triggers a decrease in FER-eGFP PM fluorescence.

Supplemental Figure S4. The increase in fluorescently-labeled intracellular punctae in response to 150 mM NaCl is not specific to FER.

Supplemental Figure S5. pH responses to root bending and RALF1 treatment.

Supplemental Figure S6. Root hair and root growth phenotypes of Arabidopsis *llg1-1* mutants.

Supplemental Figure S7. Root and hypocotyl phenotypes of Arabidopsis *llg1* mutants.

Supplemental Figure S8. Arabidopsis *llg1* mutants exhibit root growth phenotypes consistent with defects in mechanical signaling.

Supplemental Figure S9. FER-eGFP shows normal subcellular localization in *llg1-2* mutants.

Supplemental Figure S10. Cysteine protease inhibitors block the bending-induced reduction in FER-eGFP PM fluorescence.

Supplemental Figure S11. Cysteine protease inhibitors ALLN and MI-2 block wounding-induced, but not RALF1-induced, reduction in FER-eGFP PM fluorescence.

Supplemental Figure S12. Representative Western blot and quantification of band density.

## Acknowledgements

We are grateful to Dr. Marisa Otegui (University of Wisconsin, Madison), Dr. Antje Heese (University of Missouri), and Dr. Silke Robatzek (Ludwig Maximilian University of Munich) for generously sharing seeds of Arabidopsis mutants. We also gratefully acknowledge the Arabidopsis Biological Resource Center at the Ohio State University for providing insertional mutants. We would further like to acknowledge Dr. Antje Heese for guidance on how to quantify endosome formation and both Dr. Antje Heese and Linhan Sun (Pennsylvania State University) for assistance with the Western blotting protocol. The authors would further like to thank Drs. Edgar Spalding and Nathan Miller (University of Wisconsin, Madison) for making the kinematics analysis software Image Processing Toolkit v10 available. Finally, we would like to thank Drs. Aditi Bhat and Cody DePew for helpful discussions. This study was supported by NSF grant MCB1817934 and NASA grant NNX13AM47G (to G.B.M).

## Contributions

C.S.C., H.-W.S and G.B.M. designed, performed and analyzed experiments and wrote the manuscript.

Arabidopsis accessions used in this study: AT3G51550 (FER), AT5G56170 (LLG1), AT5G46330 (FLS2), AT3G05890 (RCI2B).

## Parsed Citations

Bassham, D.C., Laporte, M., Marty, F., Moriyasu, Y., Ohsumi, Y., Olsen, L.J., and Yoshimoto, K. (2006). Autophagy in Development and Stress Responses of Plants. Autophagy 2: 2–11.

Bastien, R., Legland, D., Martin, M., Fregosi, L., Peaucelle, A., Douady, S., Moulia, B., and Hï¿½fte, H. (2016). KymoRod: a method for automated kinematic analysis of rod-shaped plant organs. Plant Journal 88: 468–475.

Ben Khaled, S. (2016). Post-translational events control pattern recognition receptor trafficking to preserve PAMP responsiveness in plant immunity. Doctoral thesis, University of East Anglia

Ben Khaled, S., Postma, J., and Robatzek, S. (2015). AMoving View: Subcellular Trafficking Processes in Pattern Recognition Receptor–Triggered Plant Immunity. Annual Review of Phytopathology 53: 379–402.

Bhat, A., Depew, C.L., and Monshausen, G.B. (2021). High-resolution kinematic analysis of root gravitropic bending using RootPlot. Methods in Molecular Biology, in press

Bradford, M.M. (1976). ARapid and Sensitive Method for the Quantitation Microgram Quantities of Protein Utilizing the Principle of Protein-Dye Binding. Analytical Biochemistry 72: 248–254.

Chakravorty, D., Yu, Y., and Assmann, S.M. (2018). Akinase-dead version of FERONIAreceptor-like kinase has dose-dependent impacts on rosette morphology and RALF1-mediated stomatal movements. FEBS Letters 592: 3429–3437.

Chen, J. et al. (2016). FERONIAinteracts with ABI2-type phosphatases to facilitate signaling cross-talk between abscisic acid and RALF peptide in Arabidopsis. Proceedings of the National Academy of Sciences 113: E5519–E5527.

Cutler, S.R., Ehrhardt, D.W., Griffitts, J.S., and Somerville, C.R. (2000). Random GFP::cDNAfusions enable visualization of subcellular structures in cells of Arabidopsis at a high frequency. PNAS 97: 3718–3723.

Dejonghe, W. et al. (2019). Disruption of endocytosis through inhibition of clathrin heavy chain function. Nature Chemical Biology 15: 641–649.

Deslauriers, S.D. and Larsen, P.B. (2010). FERONIAis a key modulator of brassinosteroid and ethylene responsiveness in arabidopsis hypocotyls. Molecular Plant 3: 626–640.

Dettmer, J., Hong-Hermesdorf, A., Stierhof, Y.-D., and Schumacher, K. (2006). Vacuolar H+-ATPase Activity Is Required for Endocytic and Secretory Trafficking in Arabidopsis. The Plant Cell 18: 715–730.

Dong, Q., Zhang, Z., Liu, Y., Tao, L.-Z., and Liu, H. (2019). FERONIAregulates auxin-mediated lateral root development and primary root gravitropism. FEBS Letters 593: 97–106.

Duan, Q., Kita, D., Li, C., Cheung, A.Y., and Wu, H.-M. (2010). FERONIAreceptor-like kinase regulates RHO GTPase signaling of root hair development. Proceedings of the National Academy of Sciences of the United States of America 107: 17821–17826.

Dünser, K., Gupta, S., Kleine-vehn, J., Herger, A., Feraru, M.I., Ringli, C., and Kleine-Vehn, J. (2019). Extracellular matrix sensing by FERONIAand Leucine-Rich Repeat Extensins controls vacuolar expansion during cellular elongation in Arabidopsis thaliana. The EMBO Journal 38: e100353.

Earley, K.W., Haag, J.R., Pontes, O., Opper, K., Juehne, T., Song, K., and Pikaard, C.S. (2006). Gateway-compatible vectors for plant functional genomics and proteomics. The Plant Journal 45: 616–629.

Emans, N., Zimmermann, S., and Fischer, R. (2002). Uptake of a Fluorescent Marker in Plant Cells Is Sensitive to Brefeldin Aand Wortmannin. The Plant Cell 14: 71–86.

Erwig, J., Ghareeb, H., Kopischke, M., Hacke, R., Matei, A., Petutschnig, E., and Lipka, V. (2017). Chitin-induced and CHITIN ELICITOR RECEPTOR KINASE1 (CERK1) phosphorylation-dependent endocytosis of Arabidopsis thaliana LYSIN MOTIF-CONTAINING RECEPTOR-LIKE KINASE5 (LYK5). New Phytologist 215: 382–396.

Escobar-Restrepo, J.M., Huck, N., Kessler, S., Gagliardini, V., Gheyselinck, J., Yang, W.-C., and Grossniklaus, U. (2007). The FERONIA receptor-like kinase mediates male-female interactions during pollen tube reception. Science 317: 656–660.

Feng, W. et al. (2018). The FERONIAReceptor Kinase Maintains Cell-Wall Integrity during Salt Stress through Ca2+Signaling. Current Biology 28: 666-675.e5.

Fenteany, G., Standaert, R.F., Lane, W.S., Choi, S., Corey, E.J., and Schreiber, S.L. (1995). Inhibition of Proteasome Activities and Subunit-Specific Amino-Terminal Threonine Modification by Lactacystin. Science 268: 726–731.

Fernandes, M., Duplaquet, L., and Tulasne, D. (2019). Proteolytic cleavages of MET: the divide-and-conquer strategy of a receptor tyrosine kinase. BMB Reports 52: 239–249.

Galindo-Trigo, S., Blanco-Touriñán, N., Defalco, T.A., Wells, E.S., Gray, J.E., Zipfel, C., and Smith, L.M. (2020). CrRLK1L receptor-like kinases HERK 1 and ANJEA are female determinants of pollen tube reception. EMBO reports 21: e48466.

Gjetting, S.K., Mahmood, K., Shabala, L., Kristensen, A., Shabala, S., Palmgren, M., and Fuglsang, A.T. (2020). Evidence for multiple receptors mediating RALF-triggered Ca2+ signaling and proton pump inhibition. The Plant Journal 104: 433–446.

Gronnier, J., Franck, C.M., Stegmann, M., Defalco, T.A., Cifuentes, A.A., Dünser, K., Lin, W., Yang, Z., Kleine-Vehn, J., Ringli, C., and Zipfel, C. (2020). FERONIAregulates FLS2 plasma membrane nanoscale dynamics to modulate plant immune signaling. bioRxiv: 1–35.

Guo, H., Nolan, T.M., Song, G., Liu, S., Xie, Z., Chen, J., Schnable, P.S., Walley, J.W., and Yin, Y. (2018). FERONIAReceptor Kinase Contributes to Plant Immunity by Suppressing Jasmonic Acid Signaling in Arabidopsis thaliana. Current Biology 28: 3316–3324.

Hander, T. et al. (2019). Damage on plants activates Ca2+ -dependent metacaspases for release of immunomodulatory peptides. Science 363: eaar7486.

Haruta, M., Gaddameedi, V., Burch, H., Fernandez, D., and Sussman, M.R. (2018). Comparison of the effects of a kinase-dead mutation of FERONIAon ovule fertilization and root growth of Arabidopsis. FEBS Letters 592: 2395–2402.

Haruta, M., Monshausen, G.B., Gilroy, S., and Sussman, M.R. (2008). Acytoplasmic Ca2+functional assay for identifying and purifying endogenous cell signaling peptides in Arabidopsis seedlings: Identification of AtRALF1 peptide. Biochemistry 47: 6311–6321.

Haruta, M., Sabat, G., Stecker, K., Minkoff, B.B., and Sussman, M.R. (2014). APeptide Hormone and Its Receptor Protein Kinase Regulate Plant Cell Expansion. Science 343: 408–411.

Herger, A., Gupta, S., Kadler, G., Franck, C.M., Boisson-Dernier, A., and Ringli, C. (2019). LRR-extensins of vegetative tissues are a functionally conserved family of RALF1 receptors interacting with the receptor kinase FERONIA. bioRxiv: 1–36.

Höfte, H. (2015). The Yin and Yang of Cell Wall Integrity Control: Brassinosteroid and FERONIASignaling. Plant and Cell Physiology 56: 224–231.

Huang, H. (2021). Proteolytic Cleavage of Receptor Tyrosine Kinases. Biomolecules 11: 660.

Huck, N., Moore, J.M., Federer, M., and Grossniklaus, U. (2003). The Arabidopsis mutant feronia disrupts the female gametophytic control of pollen tube reception. Development 130: 2149–2159.

Huss, M., Ingenhorst, G., König, S., Gaßel, M., Dröse, S., Zeeck, A., Altendorf, K., and Wieczorek, H. (2002). Concanamycin A, the specific inhibitor of V-ATPases, binds to the VO subunit c. Journal of Biological Chemistry 277: 40544–40548.

Kliewer, A., Reinscheid, R.K., and Schulz, S. (2017). Emerging Paradigms of G Protein-Coupled Receptor Dephosphorylation. Trends in Pharmacological Sciences 38: 621–636.

Krol, E., Mentzel, T., Chinchilla, D., Boller, T., Felix, G., Kemmerling, B., Postel, S., Arents, M., Jeworutzki, E., Al-Rasheid, K.A.S., Becker, D., and Hedrich, R. (2010). Perception of the Arabidopsis danger signal peptide 1 involves the pattern recognition receptor AtPEPR1 and its close homologue AtPEPR2. Journal of Biological Chemistry 285: 13471–13479.

Latorraca, N.R., Masureel, M., Hollingsworth, S.A., Heydenreich, F.M., Suomivuori, C.-M., Brinton, C., Townshend, R.J.L., Bouvier, M., Kobilka, B.K., and Dror, R.O. (2020). How GPCR Phosphorylation Patterns Orchestrate Arrestin-Mediated Signaling. Cell 183: 1813– 1825.

Ledda, F. and Paratcha, G. (2007). Negative Regulation of Receptor Tyrosine Kinase (RTK) Signaling: ADeveloping Field. Biomarker Insights 2: 45–58.

Lee, D.H. and Goldberg, A.L. (1998). Proteasome inhibitors: valuable new tools for cell biologists. Trends in Cell Biology 8: 397–403.

Lemmon, M.A., Freed, D.M., Schlessinger, J., and Kiyatkin, A. (2016). The Dark Side of Cell Signaling: Positive Roles for Negative Regulators. Cell 164: 1172–1184.

Leslie, M.E. and Heese, A. (2017). Quantitative Analysis of Ligand-Induced Endocytosis of FLAGELLIN-SENSING 2 Using Automated Image Segmentation. In Plant Pattern Recognition Receptors: Methods and Protocols, L. Shan and P. He, eds (Springer Science+Business Media LLC), pp. 39–54.

Li, B., Lu, D., and Shan, L. (2014a). Ubiquitination of pattern recognition receptors in plant innate immunity. Molecular Plant Pathology 15: 737–746.

Li, C. et al. (2015). Glycosylphosphatidylinositol-anchored proteins as chaperones and co-receptors for FERONIAreceptor kinase signaling in Arabidopsis. eLife 4: 1–21.

Li, S., Sun, P., Williams, J.S., and Kao, T. (2014b). Identification of the self-incompatibility locus F-box protein-containing complex in Petunia inflata. Plant Reproduction 27: 31–45.

Li, X., Wang, X., Yang, Y., Li, R., He, Q., Fang, X., Luu, D.-T., Maurel, C., and Lin, J. (2011). Single-Molecule Analysis of PIP2;1 Dynamics and Partitioning Reveals Multiple Modes of Arabidopsis Plasma Membrane Aquaporin Regulation. The Plant Cell 23: 3780–3797.

Liao, D., Cao, Y., Sun, X., Espinoza, C., Nguyen, C.T., Liang, Y., and Stacey, G. (2017). Arabidopsis E3 ubiquitin ligase PLANT U-BOX13 (PUB13) regulates chitin receptor LYSIN MOTIF RECEPTOR KINASE5 (LYK5) protein abundance. New Phytologist 214: 1646–1656.

Lid, S.E., Gruis, D., Jung, R., Lorentzen, J.A., Ananiev, E., Chamberlin, M., Niu, X., Meeley, R., Nichols, S., and Olsen, O.-A. (2002). The defective kernel 1 (dek1) gene required for aleurone cell development in the endosperm of maize grains encodes a membrane protein of the calpain gene superfamily. PNAS 99: 5460–5465.

Lu, D., Lin, W., Gao, X., Wu, S., Cheng, C., Avila, J., Heese, A., Devarenne, T.P., He, P., and Shan, L. (2011). Direct Ubiquitination of Pattern Recognition Receptor FLS2 Attenuates Plant Innate Immunity. Science 332: 1439–1442.

Macho, A.P. et al. (2014). ABacterial Tyrosine Phosphatase Inhibits Plant Pattern Recognition Receptor Activation. Science 343: 1509– 1513.

Masachis, S., Segorbe, D., Turrà, D., Leon-ruiz, M., Fürst, U. Ghalid, M. El, Leonard, G., López-berges, M.S., Richards, T.A., Felix, G., and Pietro, A. Di (2016). Afungal pathogen secretes plant alkalinizing peptides to increase infection. Nature Microbiology 1: 16043.

Mbengue, M., Bourdais, G., Gervasi, F., Beck, M., Zhou, J., Spallek, T., Bartels, S., Boller, T., Ueda, T., Kuhn, H., and Robatzek, S. (2016). Clathrin-dependent endocytosis is required for immunity mediated by pattern recognition receptor kinases. Proceedings of the National Academy of Sciences 113: 11034–11039.

Ménard, L., Parker, P.J., and Kermorgant, S. (2014). Receptor tyrosine kinase c-Met controls the cytoskeleton from different endosomes via different pathways. Nature Communications 5: 1–14.

Monshausen, G.B., Bibikova, T.N., Messerli, M.A., Shi, C., and Gilroy, S. (2007). Oscillations in extracellular pH and reactive oxygen species modulate tip growth of Arabidopsis root hairs. Proceedings of the National Academy of Sciences of the United States of America 104: 20996–21001.

Monshausen, G.B., Bibikova, T.N., Weisenseel, M.H., and Gilroy, S. (2009). Ca2+ Regulates Reactive Oxygen Species Production and pH during Mechanosensing in Arabidopsis Roots. The Plant Cell 21: 2341–2356.

Monshausen, G.B., Miller, N.D., Murphy, A.S., and Gilroy, S. (2011). Dynamics of auxin-dependent Ca2+and pH signaling in root growth revealed by integrating high-resolution imaging with automated computer vision-based analysis. Plant Journal 65: 309–318.

Nagai, T., Yamada, S., Tominaga, T., Ichikawa, M., and Miyawaki, A. (2004). Expanded dynamic range of fluorescent indicators for Ca2+ by circularly permuted yellow fluorescent proteins. Proceedings of the National Academy of Sciences 101: 10554–10559.

Nühse, T.S., Stensballe, A., Jensen, O.N., and Peck, S.C. (2004). Phosphoproteomics of the Arabidopsis Plasma Membrane and a New Phosphorylation Site Database. The Plant Cell 16: 2394–2405.

Ono, Y., Saido, T.C., and Sorimachi, H. (2016). Calpain research for drug discovery: challenges and potential. Nature Reviews 15: 854– 876.

Paez Valencia, J., Goodman, K., and Otegui, M.S. (2016). Endocytosis and Endosomal Trafficking in Plants. Annual Review of Plant Biology 67: 309–335.

Perraki, A. et al. (2018). Phosphocode-dependent functional dichotomy of a common co-receptor in plant signaling. Nature 561: 248– 252.

Petutschnig, E.K. et al. (2014). Anovel Arabidopsis CHITIN ELICITOR RECEPTOR KINASE 1 (CERK1) mutant with enhanced pathogen-induced cell death and altered receptor processing. New Phytologist 204: 955–967.

Robatzek, S., Chinchilla, D., and Boller, T. (2006). Ligand-induced endocytosis of the pattern recognition receptor FLS2 in Arabidopsis. Genes and Development 20: 537–542.

Rotman, N., Rozier, F., Boavida, L., Dumas, C., Berger, F., and Faure, J.E. (2003). Female control of male gamete delivery during fertilization in Arabidopsis thaliana. Current Biology 13: 432–436.

Salamon, S. and Robatzek, S. (2006). Induced Endocytosis of the Receptor Kinase FLS2. Plant Signaling & Behavior 1: 293–295.

Schwessinger, B., Roux, M., Kadota, Y., Ntoukakis, V., Sklenar, J., Jones, A., and Zipfel, C. (2011). Phosphorylation-Dependent Differential Regulation of Plant Growth, Cell Death, and Innate Immunity by the Regulatory Receptor-Like Kinase BAK1. PLoS Genetics 7: e1002046.

Shen, Q., Bourdais, G., Pan, H., Robatzek, S., and Tang, D. (2017). Arabidopsis glycosylphosphatidylinositol-anchored protein LLG1 associates with and modulates FLS2 to regulate innate immunity. Proceedings of the National Academy of Sciences 114: 5749–5754.

Shen, W., Liu, J., and Li, J.-F. (2019). Type-II Metacaspases Mediate the Processing of Plant Elicitor Peptides in Arabidopsis. Molecular Plant 12: 1524–1533.

Shih, H.-W., Miller, N.D., Dai, C., Spalding, E.P., and Monshausen, G.B. (2014). The Receptor-like Kinase FERONIAIs Required for Mechanical Signal Transduction in Arabidopsis Seedlings. Current Biology 24: 1887–1892.

Stegmann, M., Monaghan, J., Smakowska-luzan, E., Rovenich, H., Lehner, A., Holton, N., Belkhadir, Y., and Zipfel, C. (2017). The receptor kinase FER is a RALF-regulated scaffold controlling plant immune signalling. Science 355: 287–289.

Takagi, J. and Uemura, T. (2018). Use of Brefeldin Aand Wortmannin to Dissect Post-Golgi Organelles Related to Vacuolar Transport in Arabidopsis thaliana. In Plant Vacuolar Trafficking, C. Pereira, ed (Humana Press), pp. 155–165.

Thynne, E. et al. (2017). Fungal phytopathogens encode functional homologues of plant rapid alkalinization factor (RALF) peptides. Molecular Plant Pathology 18: 811–824.

Toyota, M., Spencer, D., Sawai-Toyota, S., Jiaqi, W., Zhang, T., Koo, A.J., Howe, G.A., and Gilroy, S. (2018). Glutamate triggers long-distance calcium-based plant defense signaling. Science 361: 1112–1115.

Tran, D., Galletti, R., Neumann, E.D., Dubois, A., Sharif-Naeini, R., Geitmann, A., Frachisse, J.-M., Hamant, O., and Ingram, G.C. (2017). A mechanosensitive Ca2+ channel activity is dependent on the developmental regulator DEK1. Nature Communications 8: 1009.

Tsubuki, S., Saito, Y., Tomioka, M., Ito, H., and Kawashima, S. (1996). Differential Inhibition of Calpain and Proteasome Activities by Peptidyl Aldehydes of Di-Leucine and Tri-Leucine. Journal of Biochemistry 119: 572–576.

Wan, W.-L., Fröhlich, K., Pruitt, R.N., Nürnberger, T., and Zhang, L. (2019). Plant cell surface immune receptor complex signaling. Current Opinion in Plant Biology 50: 18–28.

Wang, C., Barry, J.K., Min, Z., Tordsen, G., Rao, A.G., and Olsen, O.-A. (2003). The Calpain Domain of the Maize DEK1 Protein Contains the Conserved Catalytic Triad and Functions as a Cysteine Proteinase. The Journal of Biological Chemistry 278: 34467–34474.

Xiao, Y., Stegmann, M., Han, Z., Defalco, T.A., Parys, K., Xu, L., Belkhadir, Y., Zipfel, C., and Chai, J. (2019). Mechanisms of RALF peptide perception by a heterotypic receptor complex. Nature 572: 270–274.

Yang, F., Kimberlin, A.N., Elowsky, C.G., Liu, Y., Gonzalez-Solis, A., Cahoon, E.B., and Alfano, J.R. (2019). APlant Immune Receptor Degraded by Selective Autophagy. Molecular Plant 12: 113–123.

Yin, J., Yi, H., Chen, X., and Wang, J. (2019). Post-Translational Modifications of Proteins Have Versatile Roles in Regulating Plant Immune Responses. International Journal of Molecular Sciences 20: 2807.

Yu, F. et al. (2012). FERONIAreceptor kinase pathway suppresses abscisic acid signaling in Arabidopsis by activating ABI2 phosphatase. Proceedings of the National Academy of Sciences 109: 14693–14698.

Yu, M. et al. (2020). The RALF1-FERONIA interaction modulates endocytosis to mediate control of root growth in Arabidopsis. Development 147: dev189902.

Yu, Y. and Assmann, S.M. (2018). Inter-relationships between the heterotrimeric Gβ subunit AGB1, the receptor-like kinase FERONIA, and RALF1 in salinity response. Plant, Cell & Environment 41: 2475–2489.

Zhang, X. et al. (2020). Nematode-Encoded RALF Peptide Mimics Facilitate Parasitism of Plants through the FERONIAReceptor Kinase. Molecular Plant 13: 1434–1454.

Zhao, C., Zayed, O., Yu, Z., Jiang, W., Zhu, P., Hsu, C., Zhang, L., Tao, W.A., Lozano-Durán, R., and Zhu, J.-K. (2018). Leucine-rich repeat extensin proteins regulate plant salt tolerance in Arabidopsis. PNAS 115: 13123–13128.

Zhou, J., Wang, P., Claus, L.A.N., Savatin, D. V, Xu, G., Wu, S., Meng, X., Russinova, E., He, P., and Shan, L. (2019). Proteolytic Processing of SERK3/BAK1 Regulates Plant Immunity, Development, and Cell Death. Plant Physiology 180: 543–558.

Zipfel, C., Kunze, G., Chinchilla, D., Caniard, A., Jones, J.D.G., Boller, T., and Felix, G. (2006). Perception of the Bacterial PAMP EF-Tu by the Receptor EFR Restricts Agrobacterium-Mediated Transformation. Cell 125: 749–760.

Zwiewka, M., Nodzyński, T., Robert, S., Vanneste, S., and Friml, J. (2015). Osmotic Stress Modulates the Balance between Exocytosis and Clathrin-Mediated Endocytosis in Arabidopsis thaliana. Molecular Plant 8: 1175–1187.

